# Visual experience instructs the organization of cortical feedback inputs to primary visual cortex

**DOI:** 10.1101/2022.10.12.511901

**Authors:** Rodrigo F. Dias, Radhika Rajan, Margarida Baeta, Tiago Marques, Leopoldo Petreanu

## Abstract

Cortical feedback (FB) projections are thought to modulate lower-order activity depending on learned expectations. However, whether FB inputs become bound to specific lower-order neurons depending on experience is unknown. We measured the effects of dark rearing and manipulations of experienced visual statistics on the retinotopic specificity of projections from the lateromedial (LM) visual area to layer 1 of the mouse primary visual cortex (V1). LM inputs were, on average, retinotopically matched with V1 neurons irrespective of visual experience. While the orientation tuning of LM axons determined the retinotopic position of the V1 neurons they innervated, this organization was absent in dark-reared mice. Restricting visual experience to a narrow range of orientations revealed that visual experience exerts an instructive role in the retinotopic organization of LM inputs in V1. Our observations support theories of hierarchical computation proposing that inputs from higher-order neurons to lower-order ones reflect learned hierarchical associations.

## Introduction

Perception is an active process that combines sensory stimuli with prior knowledge of the world (Summerfield and Egner, 2009). Prior knowledge disambiguates noisy stimuli, biases percepts, and shapes sensory representations (de Lange et al., 2018). Predictive processes require associating features of different complexity, e.g., locations must become associated with the objects that are usually found there and the objects themselves must be linked with their expected properties, such as their color and size (Bar, 2004). How the brain learns and stores these associations to make predictions is not well understood.

Visual perception emerges from a set of hierarchically organized cortical areas. While neurons in lower areas respond to simple visual features such as oriented bars, and have relatively small receptive fields, neurons in higher areas represent increasingly complex stimuli, and have larger receptive fields (Felleman and Van Essen, 1991; Siegle et al., 2021). Given their more complex and increasingly invariant tuning, neurons in higher-order areas are thought to shape sensory representations in lower areas depending on context and expectations through direct cortico-cortical feedback (FB) projections (Gilbert and Li, 2013; Pennartz et al., 2019). Thus, predictions might be constituted of an association between activity patterns in higher- and lower-order neurons mediated by FB connections (Bastos et al., 2012; Keller and Mrsic-Flogel, 2018; Lee and Mumford, 2003; Rao and Ballard, 1999). Consistent with this, there is increasing evidence that FB projections innervate primary visual cortex (V1) with specificity for cell and functional types (Federer et al., 2021; Huh et al., 2018; Ji et al., 2015; Marques et al., 2018; Siu et al., 2021; Young et al., 2021), as required for linking neurons with particular patterns of activity in higher- and lower-order areas. The organization of FB inputs from the lateromedial (LM) visual cortex, an area considered as the mouse homolog to primate V2 (Wang and Burkhalter, 2007), also reflects a relation between the functional properties of the afferent axons and their postsynaptic neurons in V1 (Marques et al., 2018). On average, LM axons in layer(L)1 of V1 (V1L1) innervate neurons with receptive fields (RFs) like their own, i.e., they are retinotopically matched. However, when innervating neurons with non-matching RFs, LM axons avoid cells with RFs displaced along the axis of the axons’ preferred orientation. Thus, synapses of LM axons in V1_L1_ reflect a relation between the orientation tuning of the higher-order presynaptic neurons and the spatial receptive field of lower-order postsynaptic ones, suggesting a learned association of their joint activity. If the organization of FB inputs does indeed reflect a learned association between the functional properties of higher- and lower-order neurons, visual experience should have an instructive role in shaping the connection of FB inputs in L1. Consistent with this hypothesis, FB projections to V1 are sparse at eye opening and undergo a protracted maturation in humans and mice (Berezovskii et al., 2011; Burkhalter, 1993; Dong et al., 2004), providing an extended time period during which visual experience could instruct their connectivity and organization. However, the role of visual experience in the organization of FB inputs remains uncharacterized.

Here, we measured how dark-rearing mice from different ages and manipulating experienced visual statistics affected the functional organization of LM inputs innervating V1_L1_. While both manipulation types preserved the retinotopic topography of the projection, its tuning-dependent fine-scale organization was affected. Restricting visual experience to a limited range of orientations showed that visual experience plays an instructive role in shaping the organization of the projection. Together, our results show that the organization of LM inputs in V1 reflects experienced visual statistics, supporting theories advocating for a role of descending inputs in shaping the activity of lower-order neurons according to learned associations.

## Results

### FB projections from LM to V1_L1_ are retinotopically matched in the absence of visual experience

To measure the role of visual experience in the organization of LM inputs in V1 we dark-reared two groups of mice. One group was dark reared from postnatal day(P)0 (DRP0). Another group of mice was housed on a regular 12-hour light-dark cycle from birth until P21 (DRP21) and dark reared afterwards. Thus, mice in this group had normal visual experience during the period when orientation selectivity is established and the first afferents from LM start arriving in V1 (P14-P21), but they were visually deprived during the critical period for ocular dominance plasticity and when the LM projection to V1_L1_ develops most of its synaptic inputs (P21-P60)(Dong et al., 2004; Espinosa and Stryker, 2012; Hooks and Chen, 2020; Hoy and Niell, 2015; Huberman et al., 2008; Rochefort et al., 2011; Trachtenberg, 2015). We measured how the functional organization of LM axons in V1 in the two dark-reared groups compared with normally-reared (NR) mice (Figure 1A-C). Two-month-old mice were injected with an adeno-associated virus (AAV) encoding for GCaMP6s (Chen et al., 2013) in LM and another for a red-calcium indicator jRGECO1a (Dana et al., 2016) in multiple V1 locations. We verified the accuracy of the LM injection using intrinsic signal imaging (Figure 1D) and in coronal histological sections of the visual cortex registered to the brain atlas (Figure 1E). These analyses confirmed that, in all the mice, the virus injection labelled a large fraction of both supragranular and subgranular neurons in LM. LM axons densely innervated V1 in L1, L5 and L6 (D’Souza et al., 2022; Marques et al., 2018; Young et al., 2021) while jRGECO1a-expressing neurons were mostly located in L2/3 and L5 (Figure 1F). Using a two-photon microscope equipped with two lasers and a piezo actuator to rapidly change the depth of the focal plane, we simultaneously recorded calcium activity from boutons of LM axons in L1 and somata of underlying L2/3 neurons during contralateral visual stimulation (Figure 1C). In each group, we sampled LM axons and V1 neurons at different retinotopic locations of monocular V1 (Figure S1). Using small moving bars inside a 10° by 10° square we measured the RFs of individual axonal boutons and L2/3 neurons and fitted them with a two dimensional (2D) Gaussian curve (Figure 1H). The RFs of LM boutons varied greatly and were often distant to those of the V1 neurons directly underneath (Figure 1H), indicating that diverse distal visual information is relayed to V1 neurons (Keller et al., 2020; Marques et al., 2018). We measured how the LM inputs’ RFs related to those of the V1 neurons beneath. While the RFs of LM inputs were highly scattered, the average RF of LM boutons matched that of the underlying V1 neurons in azimuth and elevation across monocular V1 in all three groups (Figure 1I). Thus, LM inputs to V1 remain topographic and, on average, retinotopically matched even in the absence of visual experience.

**Figure 1.**
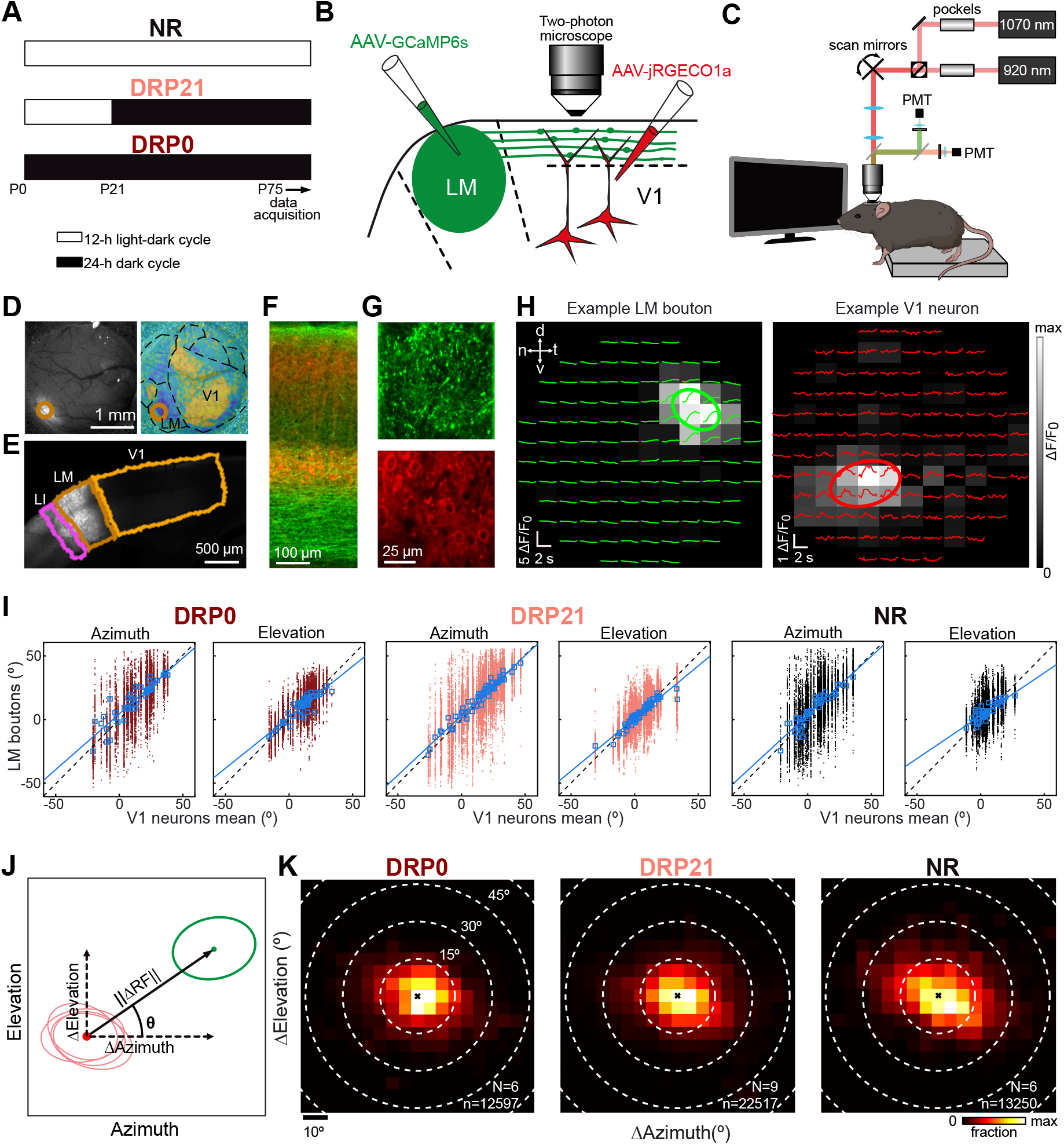
FB projections from LM to V1_L1_ are retinotopically matched in the absence of visual experience. **A**) Schematic illustration of the rearing protocols in the three groups of mice. **B**) Schematic of the dual-color imaging configuration. **C**) Schematic illustration of the experimental setup. The beams from two lasers were aligned and scanned together while independently controlling their power. **D**) Left, fiuorescent image of the cranial window showing the injection site in LM. Right, visual field sign map obtained from intrinsic signal imaging. Light orange circle indicates injection site in LM. **E**) Coronal histological section of the injection site in visual cortex. The section was registered to the mouse brain atlas to estimate area boundaries. **F**) Image of a region of V1 showing GCaMP6s-expressing axons from LM and jRGECO1a-expressing cells in V1. **G**) Two photon field of view of an example plane of GCaMP6s-expressing boutons (top) and jRGECO1a-expressing neurons (bottom). Data in panels **D-G** is from the same mouse. **H**) RF of an example LM bouton and a V1 neuron from the same session. Traces, mean fiuorescence signals for stimulus at each of the locations stimulated. Stimulus onset was at the beginning of each trace and lasted for 1.2 s. Grayscale color map, mean response 0-2 s after stimulus onset. Ellipses, fitted RFs. **I**) Visual coordinates of individual LM boutons’ RF center as a function of the mean RF centers of V1 neurons for each imaging session. Blue squares, mean values of the RF centers of LM boutons for each imaging location. Blue lines, linear fits of the mean values (statistics of linear fits in azimuth and elevation; DRP0: t_43_ =14.1, p=9e^-18^, t_43_ =13.4, p=6e^-17^, n=45 imaging locations, N=6 mice; DRP21: t_60_ =36.7, p=8e^-43^, t_60_ =24.1, p=1e^-32^, n=62, N=9; NR: t_60_ =14.9, p=7e^-18^, t_40_ =10.5, p=5e^-13^, n=42, N=6). Identity line is dashed. **J**) The relative retinotopic position (ΔRF) was defined as the difierence between the bouton’s RF center (green dot) and the mean RF center of all the V1 neurons for that imaging session (red dot). is the length of the ΔRF vector and θ is the deviation angle. **K**) 2D histogram of the distribution of ΔRF for boutons across the three visual experience groups. DRP0; n=12597 boutons, N= 6 mice; DRP21, n=22517 boutons, N=9 mice; NR, n=13250 boutons, N=6 mice. The x indicates the origin. White circles correspond to difierent distances from the origin.

For each bouton for which we could measure a RF, we computed the difference in visual coordinates between its RF center and the average RF center of all V1 neurons within the two-photon field of view (100 x 100µm^2^) (Figure 1J). This difference defines the displacement vector ΔRF, with components ΔAzimuth and ΔElevation, angle θ and length ‖ΔRF‖. As neurons in V1 are retinotopically organized, ΔRF measures the spatial relation between the RFs of the imaged boutons with their postsynaptic neurons, despite these not being identified in our recordings. To quantify and visualize the distal information relayed from LM to V1 neurons we plotted the 2D probability distribution of ΔRF for the three groups (Figure 1K). In all groups, the ΔRF probability distribution peaked around the origin, indicating that the most abundant RFs of LM inputs were retinotopically matched with those of the V1 neurons. Nevertheless, a large fraction of LM inputs had RFs distant from those of the V1 neurons although the mean ‖ΔRF‖ did not differ across experimental groups (Figure S2, one-way ANOVA: F_2,18_=1.03, p=0.38, mean ∥ Δ*RF* ∥: DRP0, 13.4±2.1°; DRP21, 14±2.4°; NR, 15.4±2.1°).

We conclude that visual experience is not required for the topographic innervation of LM inputs to retinotopically matched locations in V1_L1_ and that both retinotopically aligned and mis-aligned inputs innervate V1 neurons regardless of visual experience.

### In the absence of visual experience, LM inputs in V1_L1_ are organized based on their spatiotemporal tuning

In addition to mapping their RFs, we also measured the tuning of LM axons in V1_L1_ to moving gratings. We presented drifting gratings of 0.04 cycles per degree (c.p.d.) and 1 Hz spatial and temporal frequency, respectively, as they are known to drive a large fraction of both LM and V1 neurons (Glickfeld et al., 2013; Han et al., 2022; Marshel et al., 2011; Murakami et al., 2017). Gratings moved in 8 different directions and had 4 different orientations. From the boutons for which we could measure RFs, 63% (30587/48364 boutons) responded to these gratings (Standard Grating responsive boutons, StdGr+) (example in Figure 2A left).

**Figure 2.**
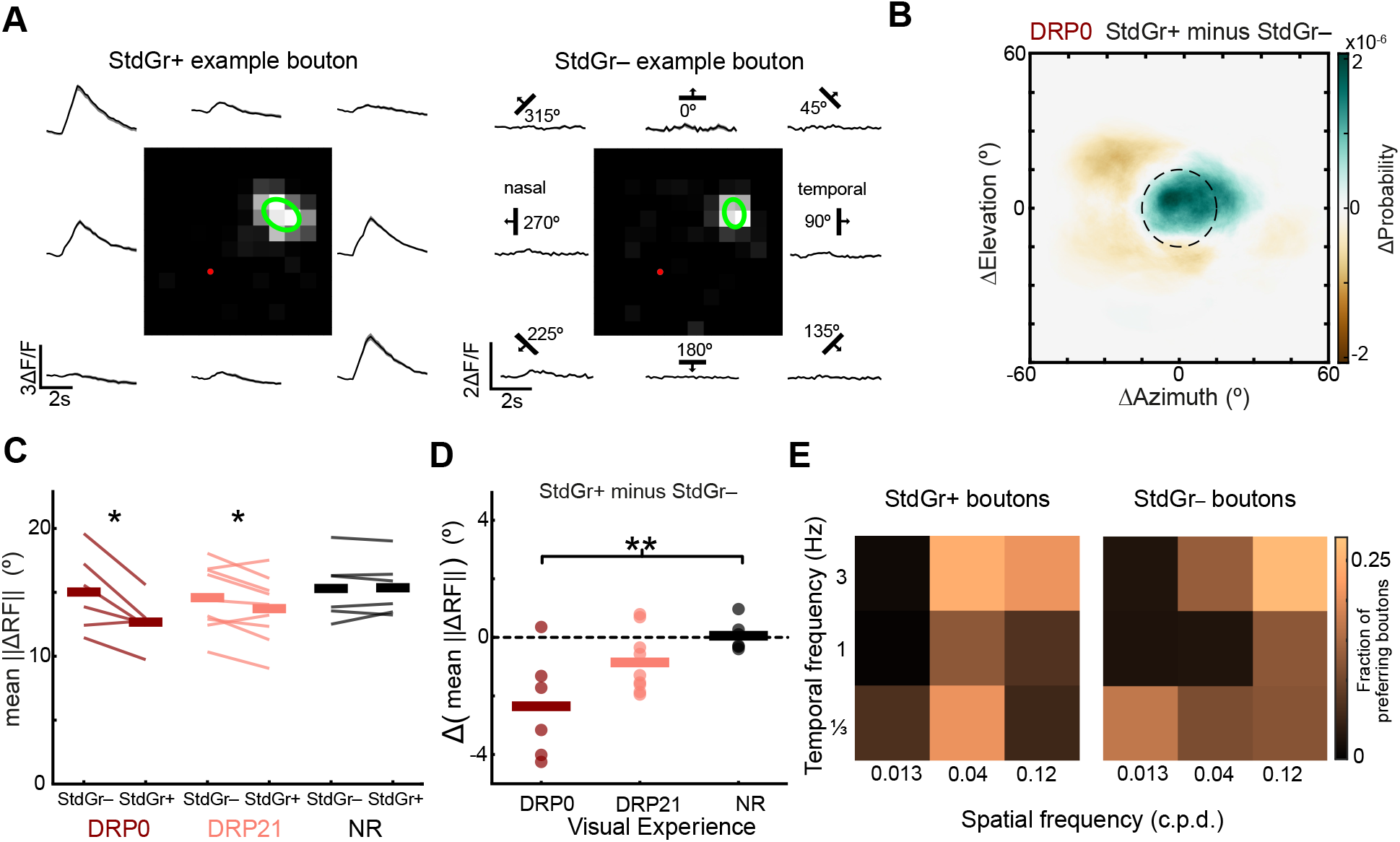
Visual experience afiects the spatiotemporal tuning-dependent organization of LM inputs in V1. **A**) RF of example Standard grating responsive (StdGr+) and Standard grating non-responsive (StdGr–) boutons. Grayscale, colormap mean response 0-2 s after stimulus onset. Green ellipse, fitted RF; red dot, mean RF center of the V1 neurons for that imaging session. Traces surrounding the RF show mean traces of the bouton to standard drifting gratings (0.04 c.p.d., 1 Hz) moving in 8 difierent directions. Shading, SEM. **B**) Colormap showing the difierence in visual coverage probability between StdGr+ and StdGr– boutons in DRP0 animals (difierence was calculated per animal before averaging, Methods). Positive values indicate a larger probability of a StdGr+ LM bouton RF to include that point in visual space in ΔRF coordinates than a StdGr– bouton. Dashed circle has a 15° radius and is centered on the origin. (StdGr+ n=8850, StdGr– n=4047 boutons, N=6 mice) **C**) The mean for StdGr+ and StdGr– boutons across the three groups DRP0, DRP21, NR. Each line indicates data from a single animal (paired t-test: DRP0, t_5_=3.24, p=0.023, N=6; DRP21, t_8_=2.44, p=0.041, N=9; NR, t_5_=-0.26, p=0.8, N=6). **D**) The difierence of the mean between the StdGr+ and StdGr– boutons across the three visual experience groups. Each dot corresponds to a single animal. One-way ANOVA F_2,18_=6.2, p=0.009; Tukey-Kramer test DRP0 vs NR, p=0.007. **E**) Fraction of StdGr+ and StdGr– boutons preferring each spatio-temporal frequency combination (StdGr+ n=2856, StdGr– n=2489 boutons, from 2 NR animals).

We then compared the spatial information relayed by StdGr+ boutons to those that had a measurable RF but did not respond to the 0.04 c.p.d. / 1 Hz gratings (Figure 2A, right, Standard gratings non-responsive boutons, StdGr–). In DRP0 mice, StdGr+ boutons were more retinotopically matched than StdGr– ones (Figure 2B,C), with StdGr+ boutons showing, on average, a lower ‖ΔRF‖ than StdGr– boutons. This difference diminished with visual experience as the ‖ΔRF‖ of the StdGr+ boutons increased, becoming indistinguishable in NR mice (Figure 2D). Thus, in the absence of visual experience, StdGr+ inputs are more retinotopically matched than StdGr– ones, but their topographic precision degrades with visual experience.

We tested whether StdGr– boutons, while not responding to the 0.04 c.p.d. / 1 Hz gratings, were visually driven by gratings of other spatiotemporal frequencies. In a separate group of mice, we presented gratings of several spatiotemporal frequencies in addition to the standard 0.04 c.p.d. / 1 Hz gratings. As expected, a fraction (73%, 1810/2489) of the StdGr– boutons responded to gratings of other frequencies, mainly preferring higher spatial and temporal frequencies (Figure 2E). Thus, StdGr+ and StdGr– boutons correspond to LM inputs with different spatiotemporal tuning (χ^2^ test: spatial frequency preference χ^2^stat=540, p=1e^-117^; temporal frequency preference χ^2^stat=25, p=3e^-6^; n=2856 StdGr+ boutons and n=2489 StdGr– boutons).

We conclude that, in the absence of visual experience, LM inputs in V1 are organized according to their spatiotemporal tuning characteristics, and that visual experience results in an increase in the fraction of StdGr+ inputs conveying distal information to V1 neurons relative to the StdGr– inputs.

### The orientation tuning-dependent organization of FB inputs in V1 requires visual experience

We then investigated whether the orientation tuning-dependent organization of LM inputs to V1 (Marques et al., 2018) was affected by the different visual deprivation protocols. The fraction of orientation-selective (OS) StdGr+ boutons (Example in Figure 2A left) was similar across the 3 groups of mice (ANOVA, F_2,18_=3.2, p=0.07), averaging 80±2% of the StdGr+ boutons. We divided the OS boutons into four groups according to their preferred orientation (Figure 3A). LM boutons preferring vertically-oriented gratings were more abundant than those tuned to other orientations (Figure 3B). We found that dark rearing did not affect this bias, as both the DRP0 and DRP21 mice show a similar overrepresentation of boutons preferring vertical orientations (Figure 3B), and visual experience did not affect the average preferred orientation of LM boutons (Watson-Williams test F_18_=2.17, p=0.14). However, visual experience beyond the first week after eye opening resulted in an increase in their orientation selectivity (Figure 3C).

**Figure 3.**
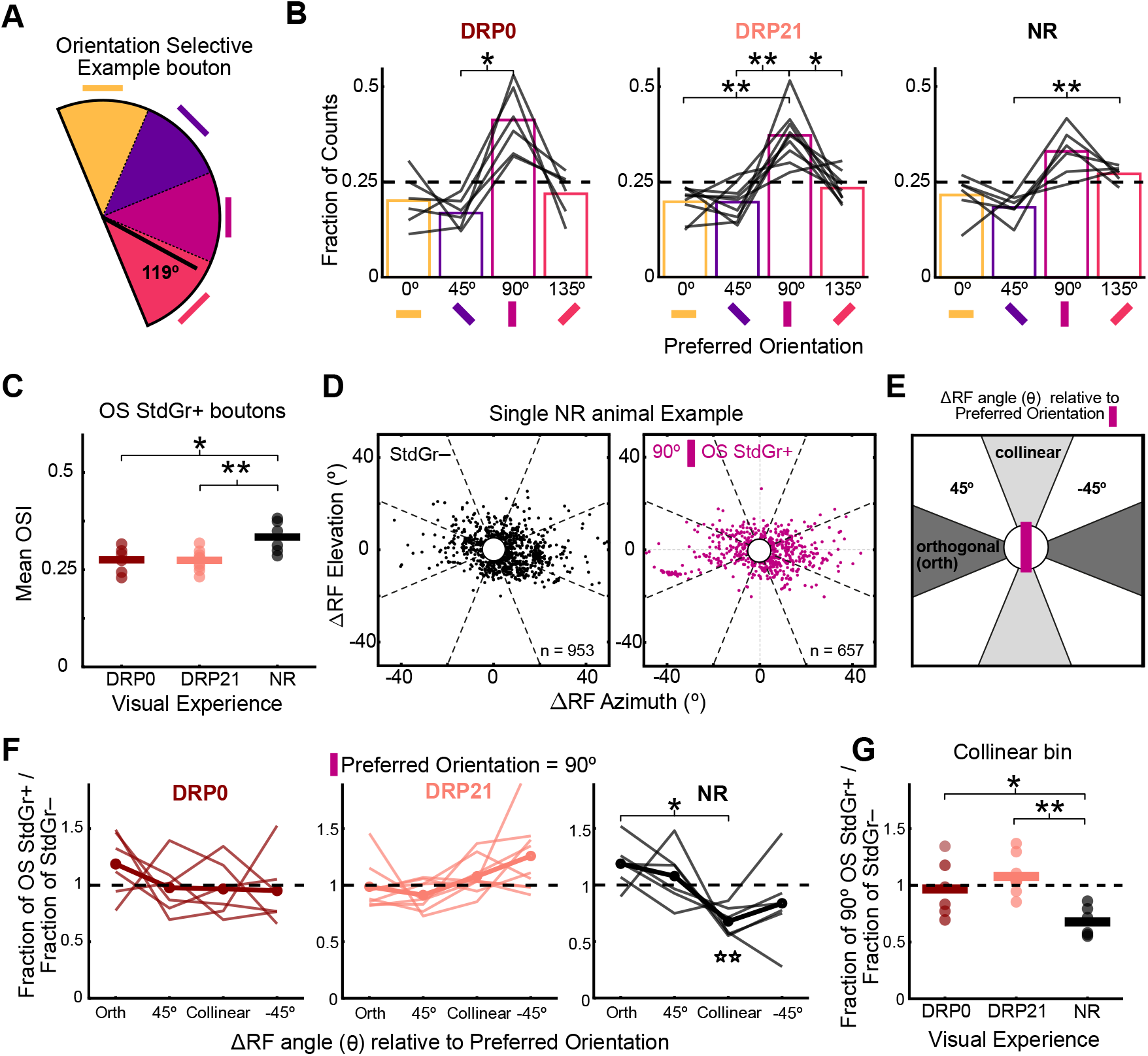
Visual experience is required for the orientation tuning-dependent organization of FB inputs in V1. **A**) StdGr+ boutons were binned in 4 groups according to their preferred orientation. Black bar, preferred orientation of the example in Figure 2**A**. **B**) Fraction of orientation selective boutons across the three visual experience groups. Only boutons with a measured RF are shown. The lines correspond to values from a single animal. (one-way Repeated Measures ANOVA: DRP0, F_5,3_=7.7, p=0.003; DRP21, F_8,3_=16.6, p=4.7e-6; NR, F_5,3_=4.6, p=0.018; Tukey-Kramer test: DRP0, 90° vs. 45° p=0.017; DRP21, 90° vs 0° p=0.002, 90° vs. 45° p=0.003, 90° vs. 135° p=0.025; NR. 45° vs. 135° p=0.002). **C**) Mean orientation selectivity index of all OS StdGr+ boutons across the three visual experience groups. (one-way ANOVA, F_2,18_=6.99, p= 0.0057; Tukey-Kramer test: DRP0 vs. NR p=0.016, DRP21 vs. NR p=0.008) **D**) ΔRF values of StdGr– and 90°-preferring OS StdGr+ boutons from a single example animal. Dashed lines, angular bins used to calculate the distribution of the deviation angle θ. Boutons with <5° were excluded. **E**) The preferred orientation of each bouton defines collinear, orthogonal, and oblique bowtie angular bins for its ΔRF deviation angle θ. **F**) Fraction of 90°-preferring boutons in each bowtie bin, normalized to the StdGr– values, across the three visual experience groups. Thin line, individual animal, thick line, mean (one-way Repeated Measures ANOVA: DRP0: F_5,3_=0.76, p=0.53; DRP21, F_8,3_=3, p=0.051; NR: F_5,3_=3.7, p=0.035; *Tukey-Kramer test Collinear vs Orthogonal bins, p=0.016; fifi, t-test of 90° tuned StdGr+ vs. StdGr– for the collinear bin, t_5_=6.01, p=0.007 after Bonferroni correction). **G**) Fraction of 90° tuned boutons deviating collinearly relative to the StdGr– ones for the three visual experience groups (one-way ANOVA for visual experience, F2,18=8.7, p=0.0023; Tukey-Kramer test: DRP0 vs NR, p=0.036; DRP21 vs NR, p=0.002).

We then asked whether visual experience was required for the orientation tuning-dependent biases of LM inputs in V1_L1_ (Marques et al., 2018). For each OS population, we measured the distribution of the ΔRF deviation angle θ (Figure 1J) using angular bins (Figure 3D,E and Figure S3A). To account for animal specific biases due to variability in the precise location of the AAV injection in LM, we normalized these angular distributions to those of the boutons that had RFs but were not responsive to these gratings (StdGr–). In these normalized plots (Figure 3F), values under 1 indicate a lower fraction of OS boutons with ΔRFs in a given angular bin relative to StdGr– boutons, while values above 1 indicate a higher fraction. The angular bins defined spaces that were orthogonal and collinear to the bouton’s preferred orientation (Figure 3E). We found that in NR mice, when taking into account all OS groups, LM boutons were less likely to deviate collinearly to the angle of their preferred orientation, as described before (Marques et al., 2018). By contrast, the ΔRF angular distribution of both DRP0 and DRP21 mice did not show any bias relative to their preferred orientation (Figure S3). When comparing across the 3 groups of mice, visual experience only affected how OS boutons displaced along the collinear bin but not along other axes relative to their preferred orientation (Figure S3). Thus, the tuning-dependent-organization of LM inputs in V1_L1_ is absent in dark-reared mice.

Next, we determined if axons with a given preferred orientation were less likely to be displaced in the collinear bin than others. In NR mice, the depletion of LM inputs in the collinear bin was robust and consistent for axons tuned to the vertical orientation (90°) but not for those tuned to other orientations (Figure 3F and Figure S4). While only vertically-tuned boutons showed a depletion of inputs deviating vertically, axons tuned to other orientations did not show a similar depletion in the vertical bin, or any other (Figure S4). Thus, the depletion along the collinear (vertical) bin is specific for the 90° tuned inputs. The depletion of 90° tuned inputs with collinearly displaced RFs was absent in both DRP0 and DRP21 groups, and different from NR (Figure 3G). Thus, visual experience affects the distribution of LM inputs in V1_L1_ in a tuning-dependent way, resulting in consistent depletion of vertically tuned axons contacting V1 neurons with vertically shifted RFs.

We conclude that while orientation preference of LM inputs in V1_L1_ is mostly unaffected by visual experience, visual experience beyond the first week after eye opening is required for their orientation tuning-dependent organization.

### Restricting visual experience to a narrow range of orientations affects the orientation tuning and RFs of LM inputs to V1_L1_

The previous dark rearing experiments showed that visual experience shapes the retinotopic specificity of LM inputs in V1_L1_ depending on their tuning to orientation (Figure 3) and spatiotemporal frequency (Figure 2). However, it is not clear from these manipulations if features of the experienced visual inputs determine the retinotopic positions innervated by LM inputs. One possibility is that visual experience is limited to a permissive role. In this case, while visual experience results in a rearrangement of LM inputs in V1_L1_, their organization would not relate to features of the experienced visual statistics. Alternatively, the rearrangements in the retinotopic specificity of LM inputs in V1_L1_ might be instructed by visual experience, such that the ΔRF distribution reflects features of the experienced stimuli. In this case, the experience-dependent depletion of vertically tuned LM inputs deviating collinearly would reflect some statistical feature of the vertical component of normal visual experience that is absent during dark rearing.

To test whether visual experience plays an instructive role in how LM inputs target V1 neurons, we measured the spatial information relayed by LM to V1_L1_ in mice raised wearing goggles with cylindrical lenses (Kreile et al., 2011; Tanaka et al., 2007). Cylindrical lenses, like stripe rearing, (Blakemore and Cooper, 1970; Sengpiel et al., 1999) restrict visual experience to a narrow range of orientations and allow to determine the extent of vision-dependent changes in neural circuits that are specific for the neurons tuned to the experienced orientations (Hirsch and Spinelli, 1970; Kreile et al., 2011; Stryker et al., 1978; Yoshida et al., 2012) (Figure 4A,B).

**Figure 4.**
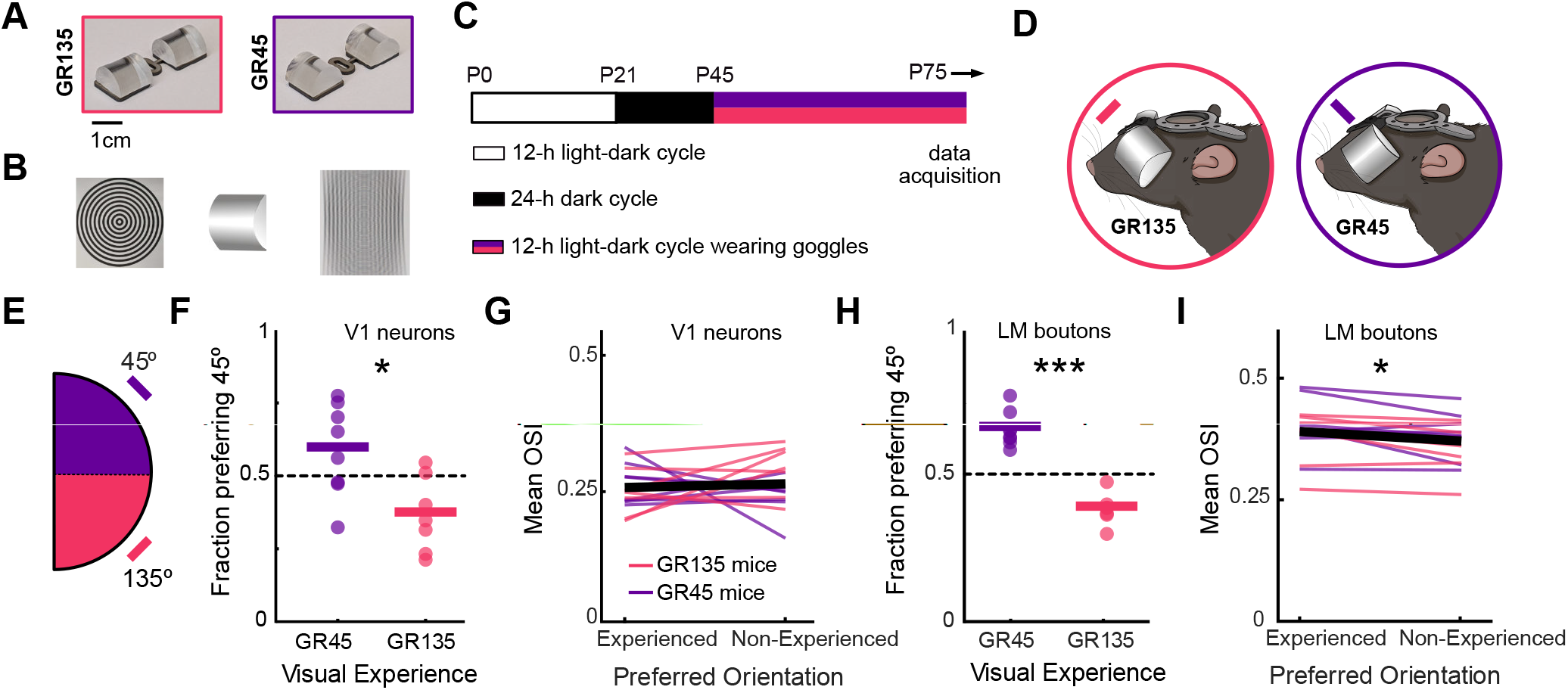
Restricting visual experience to a limited range of orientations afiects the orientation preference of V1 neurons and LM inputs. **A**) Photograph of unmounted googles used in the GR135 and GR45 groups. **B**) Concentric circles imaged with a cell phone camera with (right) and without (left) a horizontally oriented cylindrical lens in front. **C**) Schematic illustration of the goggle rearing protocol. **D**) Illustration of mice wearing 135° goggles in the GR135 group and 45° goggles in the GR45 group. **E**) OS StdGr+ boutons and V1 neurons were binned in two groups according to their preferred orientation. **F**) Fraction of V1 neurons that prefer 45° oriented gratings across the two goggle-reared groups (t-test, t_13_=3, p=0.011; GR45, N=8 mice; GR135, N=7). **G**) Mean OSI of V1 neurons tuned to the experienced and non-experienced orientations. Black line, mean; colored lines, single animals; pink, GR135 mice; violet, GR45 mice (paired t-test, t_14_=-0.4, p =0.71; N=15). **H**) Fraction of LM boutons preferring 45° oriented gratings across the two goggle-reared groups (t-test, t_13_=9.3, p=4.3 e-7; GR45, N=8 mice; GR135, N=7). **I)** Mean OSI of LM boutons tuned to the experienced and non-experienced orientations (paired t-test, t_14_=2.9, p=0.012; N=15).

Mice were first reared under normal lighting conditions until P21 without goggles, to allow for the maturation of orientation selectivity and an increased innervation of V1 neurons by StdGr+ axons with distal RFs (Figure 2C,D). Subsequently, animals were dark-reared until P45, when their head growth stabilized and head-mounted goggles with cylindrical lenses could be implanted and stably worn for more than a month. Thus, besides the week after eye opening (P14-P21), all visual experience in these mice was enriched for the orientation perpendicular to the axis of the implanted cylindrical lens (Figure 4B,C). One group was implanted with goggles restricting the visual experience to orientations around 135° (Goggle-reared 135, GR135) while another group experienced orientations around 45° (GR45) (Figure 4D). For simplicity we will refer to all orientations between 135±45° and 45±45° as the experienced orientation in GR135 and GR45 mice respectively, while orientations orthogonal to these will be referred to as the non-experienced orientation.

Wearing cylindrical lenses affected the orientation tuning of V1 neurons. Consistent with previous findings (Kreile et al., 2011), the fraction of V1 neurons tuned to 45° gratings was larger in GR45 than in GR135 mice (Figure 4F). Consequently, the mean orientation of V1 neurons was different in GR45 and GR135 groups, with a bias towards the experienced orientation in each group (circular mean of preferred orientation; GR45, 71.5±45.7°; GR135, 122.5±52.2°; Watson-Williams test F_13_=6.6, p=0.023). While goggle rearing shifted the tuning of V1 neurons towards the experienced orientation, their selectivity was unaffected, with the orientation-selectivity index (OSI) being similar in neurons tuned to the experienced orientation and those tuned to the non-experienced one (Figure 4G).

We then determined whether the tuning for orientation of LM inputs in V1_L1_ was affected in mice that experienced a narrow range of orientations. Goggle rearing resulted in an enrichment of LM boutons preferring the experienced orientation (Figure 4H). There was also a shift in the population tuning curve towards the experienced orientation (circular mean of preferred orientation; GR45, 43.4±13.5°; GR135, 113.6±6.3°; Watson-Williams test F_13_=153.1, p=1.4e^-8^). Goggle rearing also resulted in LM boutons having a slightly higher orientation selectivity than those tuned to the orthogonal, non-experienced orientation (Figure 4I).

The shape of the boutons’ RF might reflect how feedforward inputs are integrated in LM depending on the statistics of the experienced stimuli. We measured the angle of the major axis of the boutons’ RF (Methods) and compared it in GR45 and GR135 mice. Positive and negative deviations relative to the horizontal would align the RF major axis closer to the 45° and 135° orientations, respectively (Inset in Figure 5A). The angle of the major axis of the RFs of V1 neurons’ was similar across the two goggle-reared groups (Figure 5A,B), but was different for LM boutons. The RFs of LM boutons were tilted towards the angle of their preferred orientation regardless of the visual experience of the mice but also showed strong tuning-independent biases towards the angle of the experienced orientation (Figure 5C). Consequently, the angle of the LM boutons’ RFs in GR135 mice were more negative than those from the GR45 group (Figure 5D). Thus, in goggle-reared mice, the LM boutons RFs tilts towards the angle of the experienced orientation regardless of their preferred orientation.

**Figure 5.**
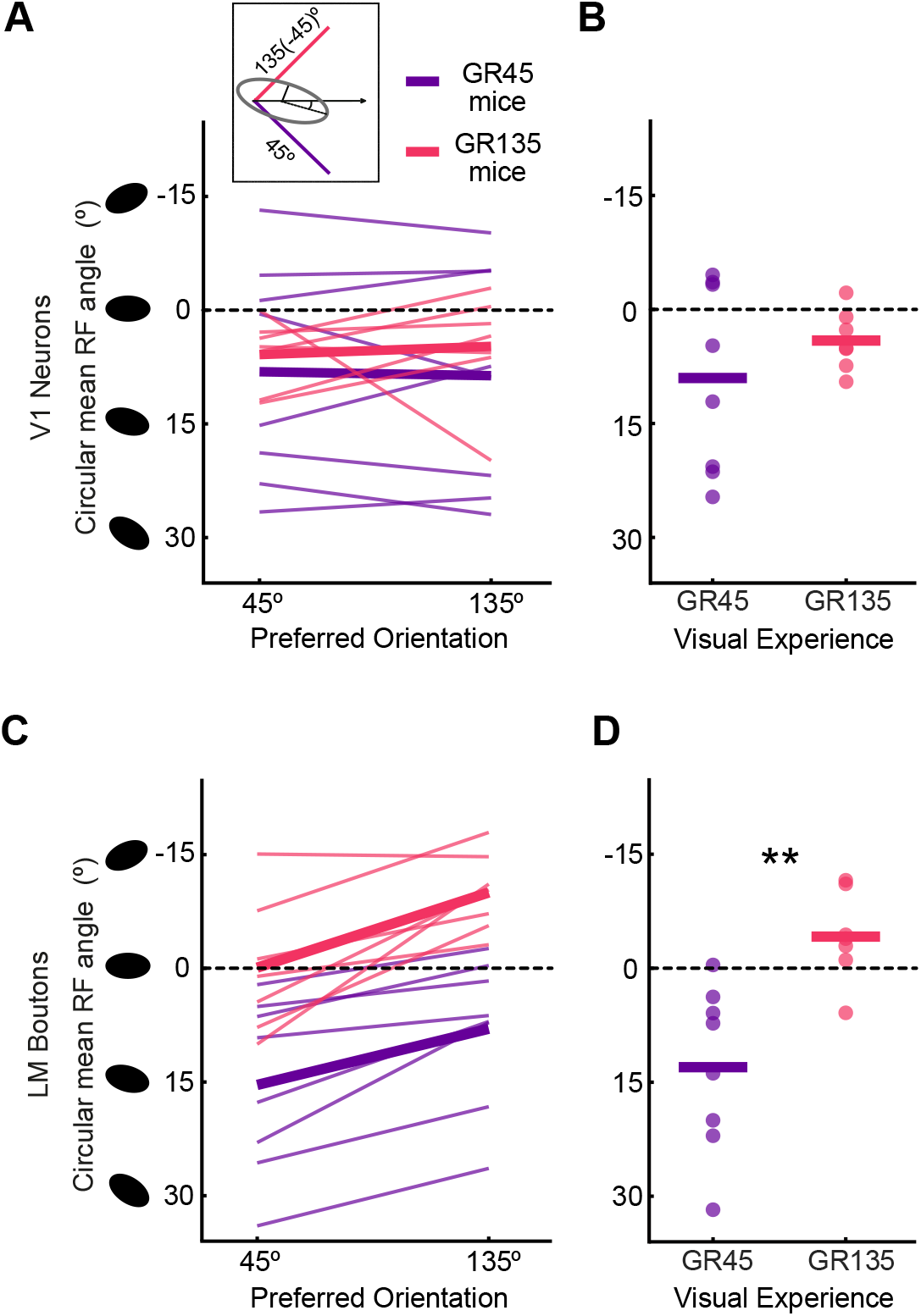
Restricting visual experience around a single orientation rearranges the receptive field shape of LM inputs but not those of V1 neurons. **A**) Circular mean of the RF long-axis angle of V1 neurons for difierent preferred orientations in GR135 (pink) and GR45 mice (violet). Thick line, average of each group; thin lines, individual animals (two-way mixed ANOVA: Visual Experience, F_1,13_=0.3, p=0.6, Preferred Orientation F_13,1_=0.02, p=0.89, Interaction F_13,1_=0.2 p=0.69; GR45, n=8 mice; GR135, n=7). Inset, the angle of the RF long axis relative the experienced orientation in GR45 (45°) and G135 mice (-45°). **B**) Circular mean of the RF long-axis angle of V1 neurons per animal for the two goggle-reared groups (Watson-Williams test, F_13_=1.03, p=0.33). **C**) Same as in **A** for LM boutons (two-way mixed ANOVA: Visual Experience F_1,13_=14, p= 0.003, Preferred Orientation F_13,1_ =33, p=6.7e^-5^, Interaction F_13,1_ =0.7, p=0.42). **D**) Same as in **C** for LM boutons (Watson-Williams test, F_13_=13.6, p= 0.0027).

We conclude that LM feedback inputs to V1 undergo large experience-dependent changes in tuning properties, resulting in an alignment of both the RFs spatial profile and orientation tuning curves towards the experienced orientation.

### Visual experience selectively organizes LM inputs tuned to experienced stimuli

The previous results showed that the tuning and RFs of LM feedback are shaped by visual experience so that they reflect the experienced orientation. We then investigated whether goggle rearing also changed how LM innervates V1_L1_. One possibility is, given that their RFs become tilted towards the angle of the experienced orientation, that LM feedback also targets neurons in V1 with RFs displaced along the angle of the experienced orientation. Such a bias would result in an overrepresentation of boutons with ΔRF angles around the experienced orientation. We found that in both GR45 and GR135 mice, LM inputs were on average retinotopically matched with V1 neurons (Figure 6A). The overall ΔRF distribution was indistinguishable between the two groups, both in the angular distribution and length of the ΔRF vectors (Figure 6B,C). In particular, there were no differences in the ‖ΔRF‖ and fraction of the LM inputs relaying information from locations deviating along the 45° or 135° angles. Thus, even though goggle rearing results in an increase in LM inputs tuned to the experienced orientation and the shape of their RFs becomes more aligned to the experienced orientation, a similar alignment along the angle of the experienced orientation is absent in their innervation of V1.

**Figure 6.**
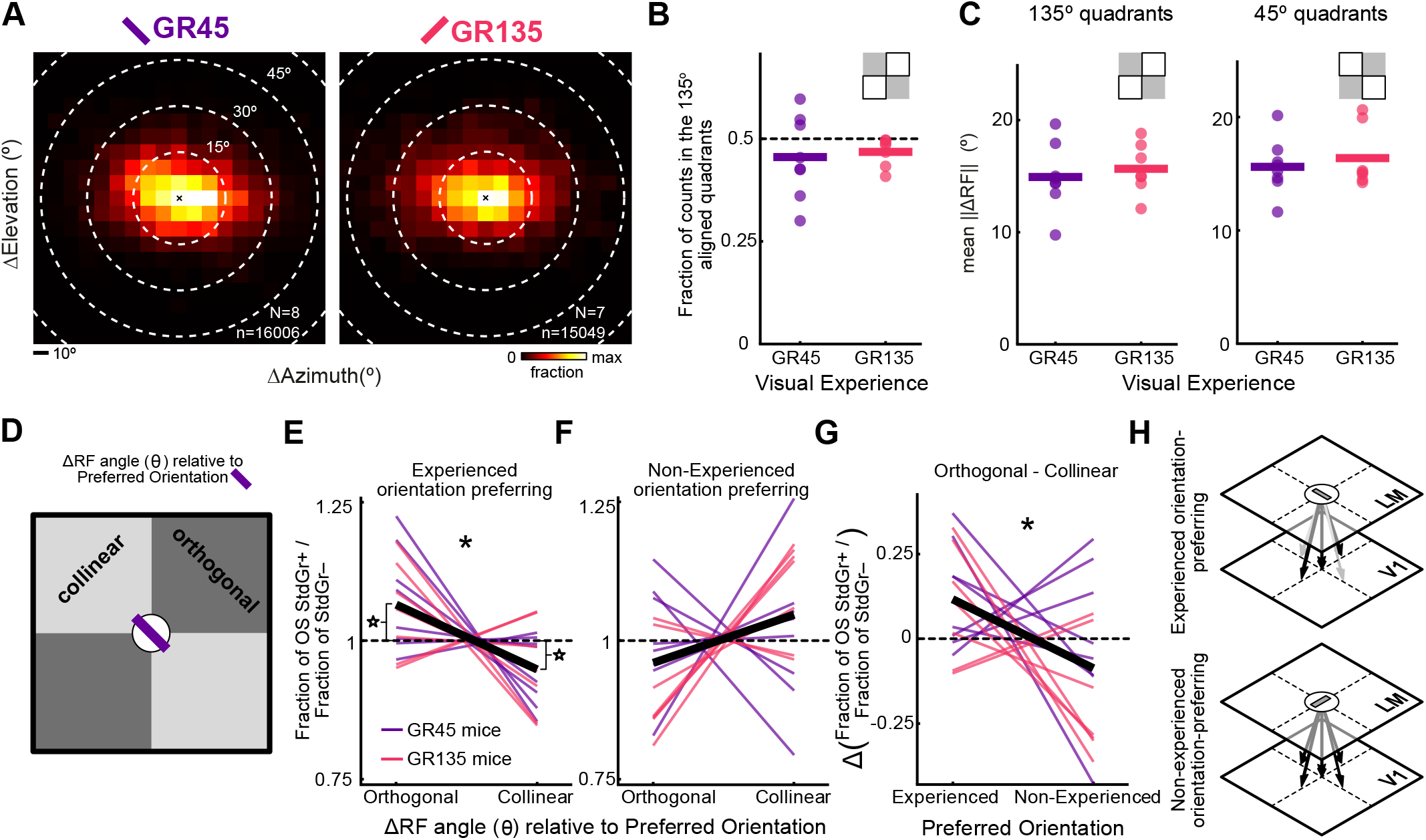
The reduction in LM inputs with ΔRF angles deviating collinearly is selective for those tuned to the experienced orientation. **A**) 2D histogram of the distribution of ΔRF for LM boutons in the two goggle groups (left) GR45 and (right) GR135 (GR45, n=16006 boutons from N=8 mice; GR135, n=15049 boutons from N=7 mice) **B**) Fraction of boutons with ΔRF in the 135° aligned quadrants (within the white area in inset) across the two goggle groups. (t-test: t_13_=-0.32, p=0.76) **C**) Mean of the boutons with ΔRF in the 135° (left) or 45° aligned quadrants (right) for the two goggle groups. (t-test: 135° quadrants, t_13_=-0.53, p=0.6; 45° quadrants, t_13_=-0.58, p=0.58). **D**) The preferred orientation of each bouton defines the collinear and the orthogonal quadrants for its ΔRF deviation angle θ. **E**) Fraction of LM boutons tuned to the experienced orientation (45° and 135° for the GR45 and GR135 groups, respectively) with ΔRF deviating in the collinear or orthogonal quadrants, normalized to the StdGr–. Black line, mean, colored lines, individual animals (pink, GR45 mice; violet, GR135 mice). (* paired t-test, t_14_=2.9, p= 0.013); fi t-test for difierence between StdGr+ and StdGr– boutons: orthogonal quadrants, t_14_=-2.9, Bonferroni corrected p=0.026; collinear quadrants, t_14_=2.8, Bonferroni corrected, p=0.028, N=15). Each line corresponds to data from a single animal. **F**) Same as in **E** for boutons tuned to the non-experienced orientations (135° and 45° for the GR45 and GR135 groups, respectively). Paired t-test, t_14_=-1.5, p= 0.15; no significant difierences between StdGr+ and StdGr–. **G**) Difierence between the fraction of StdGr+ boutons in the orthogonal and collinear bins (normalized by the StdGr– values) for the experienced and non-experienced orientations. (* paired t-test: t_14_=2.2, p= 0.042) **H**) Summary of the results in panels **E-G**. Planes represent retinotopic maps in LM and V1. Bar represents the experienced preferred orientation of the LM bouton. Only boutons tuned to the experienced orientation show a depletion of inputs to V1 neurons with RF deviating collinearly to the angle of the bouton’s preferred orientation.

However, if visual experience instructs how LM inputs innervate V1 neurons, it is expected that boutons tuned to the experienced orientation would innervate V1 neurons differently than those tuned to the non-experienced orientation. Visual experience results in orientation-tuning dependent biases in the distribution of ΔRF deviation angles (Figure 3). Thus, an instructive role of visual experience should result in boutons tuned to the experienced orientation having an organization while those tuned to the non-experienced orientation being unorganized, as observed in dark-reared mice. Consistent with this, we find that within the OS LM boutons, only those tuned to the experienced orientation (45°-preferring in GR45 and 135°-preferring in GR135), were less likely to have a ΔRF angle deviating collinearly when compared with the StdGr– ones (Figure 6D,E). In addition to the depletion of ΔRF angles deviating collinearly, these axons also show an enrichment of boutons with ΔRF angle deviating in angles perpendicular to their preferred orientation (Figure 6E), consistent with previous observations in NR mice (Marques et al., 2018). On the contrary, the ΔRF distribution of the boutons preferring the non-experienced orientation (135°-preferring in GR45 and 45°-preferring in GR135) did not show this organization and was indistinguishable from the one of StdGr– boutons (Figure 6F).

We subtracted the normalized fraction of boutons deviating collinearly from the ones deviating orthogonally in boutons tuned to the experienced and non-experienced orientations for each animal. This analysis confirmed that the angular distribution of ΔRFs was different in boutons tuned to the experienced and the non-experienced orientations (Figure 6G). Using resampling methods, we verified that these results were not due to the slight imbalance between the number of animals in the GR45 (n=8) and GR135 (n=7) groups (bootstrap test, p=0.0067).

We conclude that visual experience plays an instructive role in LM innervation of V1, resulting in a reduction of inputs to V1 neurons with RFs deviating collinearly in the angle of the axons’ preferred orientation (Figure 6H).

## Discussion

We found that the organization of LM inputs in V1_L1_ is instructed by visual experience. While visual experience did not influence the overall topography of the projection, it instructed LM inputs to innervate V1 in different retinotopic locations depending on their visual selectivity. Thus, visual experience results in FB connections that reflect a relation between the tuning of the afferent axons and the RF position of the postsynaptic neurons they innervate.

### The retinotopic targeting of LM inputs in V1_L1_ does not require visual experience

We found that LM inputs in L1 were retinotopically matched with V1 neurons even in mice that were dark reared since birth (Figure 1). Thus, like the retinal ganglion cells’ projections to the superior colliculus and the lateral geniculate nucleus (Huberman et al., 2008), visual experience is not required for their topographic organization. Unlike retinal ganglion cells, which innervate their targets before eye opening (Espinosa and Stryker, 2012; Hooks and Chen, 2020; Huberman et al., 2008), LM projections to V1_L1_ develop almost exclusively afterwards (Berezovskii et al., 2011; Dong et al., 2004). By contrast, feedforward inputs from V1 to LM approach maturation by the time of eye opening. As in other topographic projections, molecular cues could underlie the initial targeting of FB axons followed by vision-independent, activity refinement mechanisms (Espinosa and Stryker, 2012; Hooks and Chen, 2020; Huberman et al., 2008). In addition, feedforward-driven, retinotopically-matched correlated activity between V1 and higher-visual areas, as observed before eye opening (Murakami et al., 2022), could provide activity cues for organizing the targeting of FB inputs in the absence of visual inputs. While LM inputs were retinotopically matched in dark-reared mice, their precision depended on the spatiotemporal tuning of the afferents, with StdGr+ boutons being more retinotopically matched than StdGr– ones (Figure 2C). This suggests that StdGr+ afferents might have access to molecular or spontaneous activity-driven cues to guide their projections to V1 that are not available to afferents tuned to other spatiotemporal frequencies.

### The functional organization of LM feedback to V1_L1_ is instructed by experience

Experience resulted in an increase of distal information relayed to V1 neurons by StdGr+ axons relative to StdGr– ones. Thus, for StdGr+ tuned inputs, visual experience degrades the precision of the topographic map, instead of refining it, as it does with topographic projections from the retina (Hooks and Chen, 2020). This increase likely reflects a visual experience-driven learned relation between activity in LM and V1 beyond the spontaneous activity and molecular-driven cues organizing the retinotopically-matched inputs. While vision-independent mechanisms organize StdGr+ inputs in V1 with a precise topography, non-matched, non-topographic distal inputs might reflect systematic visually-driven correlations of LM and V1 neurons with distal RFs.

In agreement with this view, we found that the orientation-tuning organization of LM inputs in V1 also depends on visual experience beyond the first week after eye opening. As in previous observations (Marques et al., 2018) we found that in NR mice LM inputs were on average less likely to target V1 neurons with RFs deviating collinearly relative to feedback inputs’ preferred orientation (Figure S3). In addition, we found that this organization is present only in vertically tuned (90°) LM inputs. This tuning-dependent organization was not only dependent on visual experience but was also instructed by it (Figure 3 and Figure 6). These results show that synapses of LM inputs in V1_L1_ represent a visual experience-driven learned association between the orientation preference of LM feedback axons and the receptive field positions of their postsynaptic V1 neurons.

Orientation-selective V1 neurons are more likely to locally connect with co-oriented V1 neurons when their RFs are displaced collinearly along the preferred orientation axis (Bosking et al., 1997; Iacaruso et al., 2017; Rossi et al., 2020; Schmidt et al., 1997). If the same connectivity rules were applied to FB afferents, one would expect vertically-tuned LM inputs to be enriched, not depleted, around V1 neurons with collinearly displaced RFs. Thus, the organizing rules of LM afferents and local V1 connections might be different. One possibility is that with visual experience LM inputs are reduced in neurons that tend to be coactive with them. Such an organizing rule would be consistent with the observed depletion upon visual experience of StdGr+ inputs to V1 neurons with retinotopically matched (Figure 2) and collinearly shifted RF (Figure 3 and Figure 6).

### Implications for theories of hierarchical computation

Our results add to the recent evidence that FB inputs to visual cortex are not diffuse and unspecific, but are functionally organized and contact specific cell types (Federer et al., 2021; Huh et al., 2018; Ji et al., 2015; Marques et al., 2018; Siu et al., 2021; Young et al., 2021). They also show that at least part of this specificity i.e., its orientation-tuning specific retinotopic bias, is instructed by visual experience. These observations are consistent with theories of hierarchical computation requiring feedback inputs to learn to predict activity in lower-order neurons and selectively influence it, such as predictive coding (Bastos et al., 2012; Keller and Mrsic-Flogel, 2018; Mumford, 1992; Rao and Ballard, 1999). They are also consistent with some implementations of hierarchical learning that require learned specificity in FB connections (Lillicrap et al., 2020). Our results suggest that learned expectations about the joint LM-V1 activity might be encoded in the specificity of the LM inputs in V1_L1_. This is supported by the following observations: 1) LM connections in V1_L1_ reflect a relation between the functional properties of the higher-order presynaptic neurons (their orientation tuning) and the lower-order neurons they innervate (their spatial receptive field) and, 2) that this association depends on visual experience and is instructed by it. Thus, predictions of lower-order activity might be learnt and encoded in the connectivity of FB inputs. Once wired, the circuit might be able to distinguish previously experienced patterns of joint LM-V1 activity from novel ones, as proposed in predictive coding and other generative models of FB function. Predictive coding requires FB inputs to suppress coactive inputs (Leinweber et al., 2017; Schneider et al., 2018). This would require an enrichment of inputs to V1 neurons with RFs deviating collinearly, as collinear, co-oriented segments are enriched in natural images (Geisler, 2008; Sigman et al., 2001). Contrary to this view, we find that, with visual experience, innervation by FB inputs decreases rather than increase, in presumptively coactive V1 neurons. Thus, the way visual experience instructs LM feedback innervation in V1 does not appear to be compatible with these inputs having a role in suppressing expected lower-order activity. However, the differences in tuning properties of LM and V1 neurons to natural movies (Yu et al., 2022) suggest that their pattern of coactivation might be more complex than the one inferred from responses to oriented gratings. Therefore, we cannot discard the possibility that, while apparently contradictory to the view that LM inputs are wired for suppressing expected V1 activity, their organization still reflects targeting of coactive subpopulations of inputs during natural experience. Future experiments identifying the specific pre- and post-synaptic elements of the connections instructed by visual experience using naturalistic stimuli will provide a better understanding of the rules instructing FB connectivity and its functional implications.

## Acknowledgements

We thank Cindy Poo, Hedi Young and members of the Petreanu laboratory for comments on the manuscript, Beatriz Moura for illustrations and the Champalimaud Research Hardware Platform for technical support.

This work was supported by fellowships from Fundação para a Ciência e a Tecnologia to R.F.D. (PD/BD/114276/2016) and R.R. (PD/BD/128306/2017). T.M. was supported by the PhRMA Foundation Postdoctoral Fellowship in Informatics. L.P was supported by grants from Fundação para a Ciência e a Tecnologia (PTDC/MED-NEU/30328/2017 and PTDC/MED-NEU/6645/2020), La “Caixa” Foundation (LCF/PR/HR17/52150005) and by the Champalimaud Foundation. This work was supported by the research infrastructure CONGENTO, co-financed by Lisboa Regional Operational Programme (Lisboa2020), under the PORTUGAL 2020 Partnership Agreement, through the European Regional Development Fund (ERDF) and Fundação para a Ciência e Tecnologia (Portugal) under the project LISBOA-01-0145-FEDER-022170.

## Author contributions

R.F.D., R.R. and L.P. conceived the study. R.F.D. and R.R. performed 2-photon imaging experiments and data analysis. R.F.D., R.R. and M.B. did intrinsic signal imaging and performed surgeries. M.B. did histology. T.M. assisted with code for visual stimulation and analyses. L.P. built the dual laser 2-photon microscope. L.P., R.F.D. and R.R. wrote the paper.

## Methods Animals

All procedures were reviewed and approved by the Champalimaud Center for the Unknown Ethics Committee and performed in accordance with the Portuguese Direção Geral de Veterinária. Wild-type male C57BL/6J mice were used in all experiments. All mice had *ad libitum* access to food and water and were only used for the experiments reported here.

### Rearing Protocols

Normal-reared (NR) animals were raised on a 12-h light-dark cycle, dark-reared (DRP21) animals were raised on a 12-h light-dark cycle until weaning (P21) and then moved to a 24-h dark cycle. Fully dark-reared (DRP0) animals were born and raised on a 24-h dark cycle. The 24-h dark cycle room was kept dark and holding cages were placed behind a light-proof black curtain. Cages were taken out of the covered shelves weekly (before the craniotomy) or biweekly (after the craniotomy), one at a time under faint red light and changed (their total exposure to faint red light lasted 1-2 minutes). All mice were group housed until the day of the cranial window and headpost implantation, after which they were housed individually.

Goggle animals were raised on a 12-h light-dark cycle until weaning (P21), then moved to a 24-h dark cycle until goggle implantation (at P45) and then moved back to 12-h light-dark cycle in a cage with an enriched visual environment. The enriched visual environment consisted of a normal cage with images glued to the inner walls of the cage (3 out of the 4 walls), behind a transparent cage-shaped plastic liner. These images consisted of four sections of gratings (oriented 45° from each other, 2.35 cm wide alternating white and black bars) and two gray-scale natural images.

Goggle-reared animals were housed together with another one or more mice until the day of goggle implantation after which they were housed individually.

### Goggle design

Custom designed frames were laser cut from 0.5 mm thick stainless steel. The goggles measured 28.3 mm in length and featured two lens holding frames with an inner square aperture of 8 mm (Figure 4A). This material/thickness was robust enough that animals were not able to bend it, but flexible enough to be twisted to the intended shape for each animal’s head.

The cylindrical lenses were fabricated from 9.5 mm diameter half-round acrylic rod (www.theplasticshop.co.uk) cut in 9.5 mm square sections and glued to the two frames with Araldit epoxy (Ceys). The goggles for the GR45 and GR135 groups were identical except that the lenses were glued orthogonally in one group relative to the other one.

### Goggle implantation

Surgeries were performed when mice were 44 to 47 days old under isoflurane anesthesia (1.5-2%). Bupivacaine (0.05%, injected under the scalp) and buprenorphine (100 μl/15-30 g, 0.1 mg/kg injected subcutaneously) provided local and general analgesia. While under anesthesia, body temperature was maintained at 37 °C using a heating pad (Supertech Instruments). Sodic cefovecin, (100 μl/10g, 6 mg/kg) was administered subcutaneously to prevent infections.

Mice were anesthetized and prepared for surgery under faint red light. Subsequently, the eyes were protected and kept moist using ophthalmic ointment (Vitaminoftalmina A, Labesfal) and covered with a piece of impermeable black fabric to minimize light exposure during the surgical procedure. The skull was exposed, cleaned and protected with a thin layer of glue and dark dental cement. We also glued to the skull a M1.2 threaded stainless steel nut (Nanobeam) and partially covered it with dental cement. The frame was fixed to the implanted nut using a M1.2 screw. The frame of the goggles had a groove in which the screw could be positioned to set the anterior-posterior position of the goggles. The frames were bent so that the lenses inner sides were tangential to the eyes of the mouse while allowing grooming.

### Cranial window implantation and virus injections

When mice were 58-63 days old, they were implanted with a cranial window and on the same day injected with AAVs encoding for GCaMP6s and jRGECO1a. Surgeries were done under isoflurane anesthesia (1.5-2%). Bupivacaine (0.05%, injected under the scalp) and buprenorphine (100 μl, 0.1 mg/kg injected subcutaneously) provided local and general analgesia, dexamethasone (100 μl, 2 mg/kg, injected subcutaneously) and sodic cefovecin (100 μl/10 g, 6 mg/kg, injected subcutaneously) were used to minimize inflammation and infection, respectively. Eyes were protected and kept moist using ophthalmic ointment (Vitaminoftalmina A, Labesfal). While under anesthesia, body temperature was maintained at 37 °C using a heating pad (Supertech Instruments). A circular craniotomy of diameter 3 mm was performed over the left visual cortex (center coordinates, posterior to bregma, 3.5 mm; lateral to mid-line suture, 2.6 mm). The dura was left intact.

We targeted virus injections to visual area LM (coordinates, posterior to bregma, 3.55 mm; lateral to the mid-line suture, 3.65 mm). Virus expressing GCaMP6s (Chen et al., 2013)(AAV1-Syn-flex-GCaMP6s-WPRE-SV40, Addgene #100843) was injected (50 nL total, 10 nL/min, 250 and 550 µm deep) into the center of LM, using beveled glass pipettes (Drummond) with diameters of 8-12 μm. Pipettes were backfilled with mineral oil and frontloaded with viral suspension before injection, and virus was injected using a custom volumetric injections system based on a Narishige MO-10 manipulator. To label V1 neurons, virus expressing jRGECO1a (Dana et al., 2016) (AAV-Syn-NES-jRGECO1a-WPRE-SV40, Addgene #100854) was injected into three locations in V1 (locations correspond to the vertices of a 550 μm-wide equilateral triangle centered 3.5 mm posterior to bregma and 2.6 mm lateral to midline suture, 60 nL per injection site, 10 nL/min, 350 μm deep). (Dana et al., 2016) (AAV-Syn-NES-jRGECO1a-WPRE-SV40, Addgene #100854) was injected into three locations in V1 (locations correspond to the vertices of a 550 μm-wide equilateral triangle centered 3.5 mm posterior to bregma and 2.6 mm lateral to midline suture, 60 nL per injection site, 10 nL/min, 350 μm deep). To prevent backflow during withdrawal, the pipette was kept in the brain for over 5 min before being pulled up.

An imaging window was constructed from 3 layers of microscope cover glass (Fisher Scientific, no. 1) joined with an UV-curable optical glue (Norland). The window was embedded into the craniotomy and secured in place using black dental cement (Lang dental). A custom-designed iron headpost was attached to the skull with black dental cement.

In DRP0, DRP21 and goggle-reared animals, the animals were anesthetized and prepared for surgery under faint red light. Subsequently, the eyes were protected and kept moist using ophthalmic ointment (Vitaminoftalmina A, Labesfal) and covered with a piece of impermeable black fabric to minimize light exposure during the surgical procedure.

In goggle-reared animals dental cement was carefully removed to expose the skull before the cranial window implantation and virus injections. The goggles were detached by removing the holding screw, the eyes were covered with black fabric. Goggles were screwed back in place at the end of the surgery.

### Intrinsic signal imaging

Two weeks after cranial window implantation and before starting two-photon imaging, intrinsic signal imaging was used to obtain retinotopic maps of the primary visual cortex and surrounding visual areas in NR, GR45 and GR135 mice as described before (Marques et al., 2018). Goggles were removed before intrinsic signal imaging and reattached immediately afterwards. To minimize visual exposure for the DRP21 and DRP0 animals, intrinsic signal imaging was performed only after the 2-photon imaging experiments were completed. Mice were head-fixed and lightly anesthetized with isoflurane (1%) and injected intramuscularly with chlorprothixene (1 mg/kg). The eyes were coated with a thin layer of silicone oil (Sigma-Aldrich) to ensure optical clarity during visual stimulation. Optical images of cortical intrinsic signals were recorded using a Retiga QIClick camera (QImaging) controlled using Ephus (Suter et al., 2010) with a high magnification zoom lens (Thorlabs) focused at the brain surface under the glass window at 5 Hz. To measure intrinsic hemodynamic responses, the surface of the cortex was illuminated with a 620 nm red light-emitting diode (LED) while a drifting bar was presented to the right eye in a monitor. An image of the cortical vasculature under the window was obtained using a 535 nm green LED. After the intrinsic signal imaging session, a 430 nm blue LED was used to confirm that the GCaMP6s injection was in LM. Azimuth and elevation maps of the visual cortical areas were obtained by calculating the phase for each pixel of the discrete Fourier transform at the visual stimulation frequency (Kalatsky and Stryker, 2003). Hemodynamic delay was cancelled by subtracting the phase maps of the experiments from opposing moving stimuli.

### 2-photon imaging

We started imaging mice when they were around 75-days old and imaged them for 7.9±3.4 days (range 3-17 days). We used a custom microscope (based on the MIMMS design, Janelia Research Campus, https://www.janelia.org/open-science/mimms-10-2016) equipped with a resonant scanner. GCaMP6s was excited using a Ti:Sapphire laser (Chameleon Ultra II, Coherent) tuned at 920 nm. jRGECO1a was excited using an ultrafast fiber laser (Fidelity-2, Coherent) tuned at 1070 nm. The beams from the two lasers were combined and aligned using a polarizing beam splitter cube and their power was independently controlled using Pockels Cells (ConOptics). We used GaAsP photomultiplier tubes (10770PB-40, Hamamatsu) and a 16x (0.8 NA) objective (Nikon). We performed volumetric imaging by scanning in the axial direction with a piezo actuator (Physik Instrumente). Data from the time of the piezo flyback was discarded. To prevent light contamination from the monitor we synchronized the stimulus presentation to the flyback of the resonant scanner using a custom electronic board and modified a computer monitor to control its LED light source. The microscope was controlled using ScanImage (Pologruto et al., 2003) (Vidrio technologies). The objective optical axis was perpendicularly aligned to the imaging window. Two-photon imaging began 2 weeks after viral injections and implantation of the cranial window and continued for another 2-3 weeks.

Mice were head-fixed and anesthetized with isoflurane (0.7-1%) and injected intramuscularly with chlorprothixene (1 mg/kg). Goggles were removed and reattached at the end of the imaging session. The eyes were coated with a thin layer of silicone oil. Visual stimuli were presented to the right eye in a monitor aligned to the horizontal axis parallel to the headpost. Body temperature was maintained at 37 °C using a heating pad (Supertech Intruments). We simultaneously imaged at 4 different depths (100 × 100 µm, 512 × 512 pixels, 40 µm interplane distance, at 6 Hz). Images of FB axons were taken from the top two planes at a depth ranging from 10-80 µm from the dura. Images of L2/3 neurons were taken from the third and fourth planes, with depths ranging from 90-160 µm. Laser power in both beams averaged between 55 to 75 mW. To minimize cross talk between the different calcium indicators (Dana et al., 2016) the 1070 nm beam was turned off (<1 mW of power) when scanning in L1 and the 920 nm one was turned off when scanning in L2/3. Imaging was done over monocular primary visual cortex based on intrinsic signal imaging maps and vessels.

For DRP0 and DRP21 a quick receptive field protocol (same stimulus as below repeated only 3 times) was run to assess the retinotopic location of the neurons and decide on the imaging location.

### Visual Stimulation

Visual stimuli were presented on a LED (BenQXL2411-B 144 Hz 24 “, stimulus presented at 60 Hz refresh rate) monitor positioned 11 cm from the eye and at 30° relative to the axis of the body of the mouse to provide stimulation to the right eye. This arrangement allowed the presentation of stimuli in a portion of the visual of 120° in azimuth and elevation. Visual stimuli were generated using Matlab and Pychophysics Toolbox (Brainard, 1997).

For measuring the receptive fields of neurons and boutons, a small moving bar (8° wide, moving at 25 °/s) was presented inside a 10° by 10° square for 1.2 s (Marques et al., 2018). In each trial, the bar was presented moving sequentially in the four cardinal directions in a pseudorandom fashion. A spherical correction was applied to compensate for the monitor flatness (Marshel et al., 2011) and visual space was measured in azimuth and elevation coordinates. Stimuli were presented in an isometric, circular grid pattern, spaced 10° between locations, covering 120° in azimuth and elevation (Figure 1H). A total of 14 repetitions were presented in each grid location.

To measure orientation tuning, full screen, full contrast gratings moving in one of eight directions (cardinals and obliques) were displayed on the monitor. Gratings had a spatial frequency of 0.04 c.p.d. and a temporal frequency of 1 Hz. Each grating type was repeated for 20 times. Each trial lasted 5 s with 1 s of visual stimulation and 4 s of gray screen.

For Figure 2E we presented full-screen, full-contrast gratings moving in one of four directions (cardinals only) on the monitor with a larger range of spatial and temporal frequencies in 2 separate NR mice, while recording from LM boutons (temporal frequencies in Hz, 1/3, 1, 3; spatial frequencies in cycles per degree, 0.013,0.04, 0.12, each one repeated 20 times). Each trial lasted 4 s with 1 s of visual stimulation and 3 s of gray screen. Each trial lasted 4 s with 1 s of visual stimulation and 3 s of gray screen.

### Histology

After completion of imaging experiments, mice were deeply anesthetized, transcardially perfused, and fixed overnight with 4% paraformaldehyde in 0.1 M phosphate buffer, pH 7.4. The brain was cut into 50 μm-thick coronal sections with a vibratome (Leica). GCaMP6s-expressing cells and processes were detected using a polyclonal anti-GFP antibody (1:4000, Abcam, ab13970) and an AlexaFluor 488-conjugated secondary antibody (1:1000, Invitrogen by ThermoFisher, A-11039) and counterstained with DAPI. We imaged the visual areas at high resolution using a slide scanner with a 20x objective (Axio Scan.ZI, Zeiss). We used QuickNII and VisuAlign (Puchades et al., 2019) to align the brain sections to the Allen Mouse Brain Atlas and identify the boundaries of the visual cortical areas. Animals in which the injection site was partially in V1 or when it was mostly outside of LM were discarded. While most of the GCaMP6s-expressing neurons were confined to LM, parts of adjacent area laterointermediate (LI) area were also labelled in some mice. Thus, as LI projections to V1 are much sparser than those from LM (Morimoto et al., 2021; Wang et al., 2012) most of the boutons we recorded originated from LM neurons.

### Data Analysis

All analyses were performed using custom-written code in Matlab.

#### Preprocessing of calcium data

Imaging sessions were registered, regions of interest (ROIs) were automatically detected and fluorescence traces were extracted using Suite2p (Pachitariu et al., 2017). ROIs were then manually curated and most (98.3±0.6%, measured across 23 random imaging session across 10 animals) encompassed single boutons. Baseline fluorescence F_0_ was extracted as the 30^th^ percentile of the fluorescent trace using a 60 s sliding window and the ΔF/F_0_ was calculated as follows:

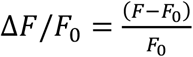

For the calculation of the neurons’ ΔF/F_0_ during the measurements of orientation tuning we corrected for neuropil contamination in Suite2p using a coefficient *r* = 0.7.

#### Measurements of receptive fields

For each ROI, responses at each location were baseline subtracted (baseline: 0.2 s before stimulus onset) and averaged across repetitions. We calculated the stimulus response *R*(*az, el*) at each location by averaging the responses during stimulus presentation (1.2 s), where *az* and *el* are the retinotopic coordinates azimuth and elevation. The array of *R*(*az, el*) responses across locations was fitted with a two-dimensional (2D) Gaussian, using the Matlab implementation of the least-squares Levenberg-Marquardt algorithm:

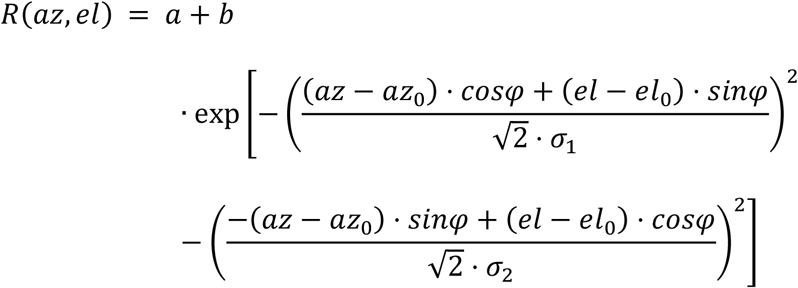

where (*az*_0_, *el*_0_) is the center of the 2D Gaussian in azimuth and elevation, σ_1_ and σ_2_ are the s.d. along the two axes of the Gaussian, *φ* is the rotation angle of the Gaussian in relation to the azimuth-elevation coordinate system, and *a* and *b* are offset and amplitude parameters. The RF boundary was considered to be the ellipse defined by the center (*az*_0_, *el*_0_) and the s.d. of the fitter 2D Gaussian (Figure 1H):

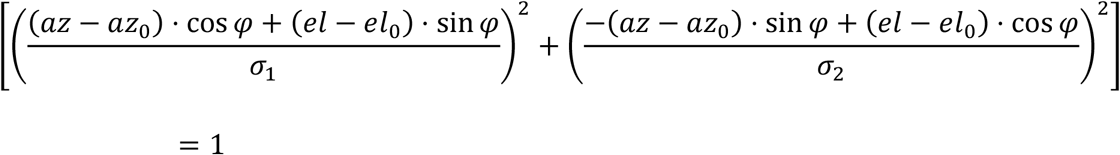

Only fits with *R*^2^ > 0.4 for boutons and *R*^2^ > 0.3 for neurons and s.d. ∈ [5,50]° and with centers within 60° from the center of the screen were considered for analysis. Lower *R*^2^ fits resulted in unreliable estimates of RF position and shape. Boutons with RFs that fulfilled the quality criteria: DRP0, 17±9%, 12597/78500; DRP21, 20±6%, 22517/111098; NR, 18±8%, 13250/76057; GR45, 14±5%, 16006/122195; GR135, 12±6%, 15049/121036.

To analyze the shape and tilt of the RFs (Figure 5) we excluded circularly shaped RF by analyzing only those with eccentricity values larger than 0.7, with eccentricity defined as:

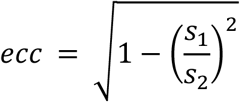

where *s*_1_ and *s*_2_ are the minor and major axis of the RF respectively.

We extracted the long and short RF axis from the 2D Gaussian fit. The angle (*α*) of the long axis was defined as 0° when it was aligned to the azimuth axis. It had positive or negative values when tilted towards lower or higher elevations, respectively (Figure 5A, inset).

To obtain the visual coverage map shown in (Figure 2B) we summed individual RFs (represented by a binarized ellipse) in ΔRF coordinates for each population. These summation maps were normalized by the total number of boutons of each population. Thus, they indicate the probability that a given position in relative visual space is represented by one converging LM bouton (Marques et al., 2018). The two maps obtained for StdGr+ and StdGr– boutons were subtracted (per animal) and averaged to get the visual coverage probability difference shown in Figure 2B.

#### Analysis of Tuning properties

For each ROI, ΔF/F_0_ traces were averaged over all repetitions for each stimulus type after subtracting the baseline fluorescence (average fluorescence 0.5 s before stimulus presentation). The mean stimulus response was obtained by averaging the mean fluorescence trace between the stimulus onset and after 0.2 s after stimulus offset. The response for each orientation was obtained by averaging the responses to two opposing directions. The signal to noise ratio (SNR) was calculated as the ratio of the maximum response (*R*_*max*_) to noise (σ).

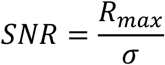

σ was measured as the mean of the standard deviation of the fluorescence trace during the baseline period (0.5 s before the stimulus). Only ROIs with an *SNR*>1 were considered to be reliably driven by the grating stimulus and included in the analysis.

ROIs were considered to be orientation selective (OS) if their responses to the preferred orientation were significantly larger than to the orthogonal orientation (p<0.05 for a t-test). Boutons were considered non-selective (StdGr–) if they were nonresponsive to any moving grating. To determine the preferred orientation, we first took the average response to the two opposing directions and then calculated the vectorial sum of the responses to the four orientations. The angle of the resulting vector was defined as the preferred orientation and the normalized amplitude is the OSI. Preferred orientation distributions were binned using 45° bins centered at the same angles as the gratings that were shown (0°, 45°, 90° and 135°, as seen in Figure 3A). When analyzing recordings from the two goggle-reared groups we used bigger bins (90°) centered at 45° and 135° (as seen in Figure 4E) to accommodate for the variability of the angle of the goggles relative to the mice’s head across animals.

We only used OS StdGr+ boutons that had a RF to obtain the preferred orientation distributions shown in Figure 3B and Figure 4H. We used all grating responsive V1 neurons for the OS distribution in Figure 4F.

#### Angular distribution of relative retinotopy

We calculated the angular distribution of the ΔRF deviation angle θ (Figure 1J) using four bowtie bins of 45° (Figure 3D,E) or two of 90° when analyzing goggle-reared mice (Figure 6D). We discarded ROIs with ‖ΔRF‖ of less than 5° due to the imprecise nature of angular measurements close to the origin of a coordinate system (Figure 3D). To correct for possible biases in the ΔRF distribution across mice, we divisively normalized the bowtie angular counts of StdGr+ boutons to those of the StdGr– boutons (Figure 3F and Figure 6E).

For the analysis shown in Figure S3B, we averaged the values across the four different OS sub-populations (Figure S3A).

### Statistical analyses

All values in text are presented as mean ± standard deviation. Statistical analyses were performed in Matlab for the tests specified using functions ttest, ttest2, anova1, crosstab, simple_mixed_anova, ranova, multcompare polyfit and corrcoef. Circular statistics (mean, variance and Watson-Williams test) were done using the CircStat toolbox (Berens, 2009). Tests were two-sided and corrected for multiple comparisons using the Bonferroni correction when indicated in the figure legend. ANOVAs were followed by Tukey-Kramer post-hoc tests.

To test whether the depletion in the collinear bin to the experienced orientation was not due to the different number of animals in the goggle-reared groups, we performed a bootstrap test equalizing the number of animals in the two groups. We sampled with replacement 7 animals from each goggle-reared group 10000 times. In each iteration we calculated for each animal the difference in the fraction of normalized counts between the orthogonal and collinear bins for boutons tuned to the experienced and the non-experienced orientations as in Figure 6G, subtracted the two, and averaged across animals. We calculated the p-value as the fraction of iterations in which the difference between the orthogonal and collinear bins for boutons tuned to experienced orientation was lower than that of those tuned to the non-experienced orientation.

## References

Bar, M. (2004). Visual objects in context. Nat. Rev. Neurosci. 5, 617–629.

Bastos, A.M., Usrey, W.M., Adams, R. a., Mangun, G.R., Fries, P., and Friston, K.J. (2012). Canonical microcircuits for predictive coding. Neuron 76, 695–711.

Berens, P. (2009). CircStat: A MATLAB Toolbox for Circular Statistics. J. Stat. Softw. 31, 128–129.

Berezovskii, V.K., Nassi, J.J., and Born, R.T. (2011). Segregation of feedforward and feedback projections in mouse visual cortex. J. Comp. Neurol. 519, 3672–3683.

Blakemore, C., and Cooper, G.F. (1970). Development of brain depends on visual environment. Nature 228, 477–478.

Bosking, W.H., Zhang, Y., Schofield, B., and Fitzpatrick, D. (1997). Orientation selectivity and the arrangement of horizontal connections in tree shrew striate cortex. J. Neurosci. 17, 2112–2127.

Brainard, D.H. (1997). The Psychophysics Toolbox. Spat. Vis. 10, 433–436.

Burkhalter, A. (1993). Development of Forward and Feedback Connections between Areas V1 and V2 of Human Visual Cortex. Cereb. Cortex 3, 476–487.

Chen, T.-W., Wardill, T.J., Sun, Y., Pulver, S.R., Renninger, S.L., Baohan, A., Schreiter, E.R., Kerr, R. a., Orger, M.B., Jayaraman, V., et al. (2013). Ultrasensitive fluorescent proteins for imaging neuronal activity. Nature 499, 295–300.

D’Souza, R.D., Wang, Q., Ji, W., Meier, A.M., Kennedy, H., Knoblauch, K., and Burkhalter, A. (2022). Hierarchical and nonhierarchical features of the mouse visual cortical network. Nat. Commun. 13, 503.

Dana, H., Mohar, B., Sun, Y., Narayan, S., Gordus, A., Hasseman, J.P., Tsegaye, G., Holt, G.T., Hu, A., Walpita, D., et al. (2016). Sensitive red protein calcium indicators for imaging neural activity. Elife 5, e12727.

Dong, H., Wang, Q., Valkova, K., Gonchar, Y., and Burkhalter, A. (2004). Experience-dependent development of feedforward and feedback circuits between lower and higher areas of mouse visual cortex. Vision Res. 44, 3389–3400.

Espinosa, J.S., and Stryker, M.P. (2012). Development and Plasticity of the Primary Visual Cortex. Neuron 75, 230–249.

Federer, F., Ta’afua, S., Merlin, S., Hassanpour, M.S., and Angelucci, A. (2021). Stream-specific feedback inputs to the primate primary visual cortex. Nat. Commun. 12, 228.

Felleman, D.J., and Van Essen, D.C. (1991). Distributed hierarchical processing in the primate cerebral cortex. Cereb. Cortex 1, 1–47.

Geisler, W.S. (2008). Visual Perception and the Statistical Properties of Natural Scenes. Annu. Rev. Psychol. 59, 167–192.

Gilbert, C.D., and Li, W. (2013). Top-down influences on visual processing. Nat. Rev. Neurosci. 14, 350–363.

Glickfeld, L.L., Andermann, M.L., Bonin, V., and Reid, R.C. (2013). Cortico-cortical projections in mouse visual cortex are functionally target specific. Nat. Neurosci. 16, 219–226.

Han, X., Vermaercke, B., and Bonin, V. (2022). Diversity of spatiotemporal coding reveals specialized visual processing streams in the mouse cortex. Nat. Commun. 13, 3249.

Hirsch, H.V.B., and Spinelli, D.N. (1970). Visual Experience Modifies Distribution of Horizontally and Vertically Oriented Receptive Fields in Cats. Science (80-.). 168, 869–871.

Hooks, B.M., and Chen, C. (2020). Circuitry Underlying Experience-Dependent Plasticity in the Mouse Visual System. Neuron 106, 21–36.

Hoy, J.L., and Niell, C.M. (2015). Layer-Specific Refinement of Visual Cortex Function after Eye Opening in the Awake Mouse. J. Neurosci. 35, 3370–3383.

Huberman, A.D., Feller, M.B., and Chapman, B. (2008). Mechanisms Underlying Development of Visual Maps and Receptive Fields. Annu. Rev. Neurosci. 31, 479–509.

Huh, C.Y.L., Peach, J.P., Bennett, C., Vega, R.M., and Hestrin, S. (2018). Feature-Specific Organization of Feedback Pathways in Mouse Visual Cortex. Curr. Biol. 28, 114-120.e5.

Iacaruso, M.F., Gasler, I.T., and Hofer, S.B. (2017). Synaptic organization of visual space in primary visual cortex. Nature 547, 449–452.

Ji, W., Gamanut, R., Bista, P., D’Souza, R.D., Wang, Q., and Burkhalter, A. (2015). Modularity in the Organization of Mouse Primary Visual Cortex. Neuron 87, 632–643.

Kalatsky, V. a, and Stryker, M.P. (2003). New paradigm for optical imaging: temporally encoded maps of intrinsic signal. Neuron 38, 529–545.

Keller, G.B., and Mrsic-Flogel, T.D. (2018). Predictive Processing: A Canonical Cortical Computation. Neuron 100, 424–435.

Keller, A.J., Roth, M.M., and Scanziani, M. (2020). Feedback generates a second receptive field in neurons of the visual cortex. Nature 582, 545–549.

Kreile, A.K., Bonhoeffer, T., and Hübener, M. (2011). Altered visual experience induces instructive changes of orientation preference in mouse visual cortex. J. Neurosci. 31, 13911–13920.

de Lange, F.P., Heilbron, M., and Kok, P. (2018). How Do Expectations Shape Perception? Trends Cogn. Sci. 22, 764–779.

Lee, T.S., and Mumford, D. (2003). Hierarchical Bayesian inference in the visual cortex. J. Opt. Soc. Am. A. Opt. Image Sci. Vis. 20, 1434–1448.

Leinweber, M., Ward, D.R., Sobczak, J.M., Attinger, A., and Keller, G.B. (2017). A Sensorimotor Circuit in Mouse Cortex for Visual Flow Predictions. Neuron 95, 1420-1432.e5.

Lillicrap, T.P., Santoro, A., Marris, L., Akerman, C.J., and Hinton, G. (2020). Backpropagation and the brain. Nat. Rev. Neurosci. 21, 335–346.

Marques, T., Nguyen, J., Fioreze, G., and Petreanu, L. (2018). The functional organization of cortical feedback inputs to primary visual cortex. Nat. Neurosci. 21, 757–764.

Marshel, J.H., Garrett, M.E., Nauhaus, I., and Callaway, E.M. (2011). Functional specialization of seven mouse visual cortical areas. Neuron 72, 1040–1054.

Morimoto, M.M., Uchishiba, E., and Saleem, A.B. (2021). Organization of feedback projections to mouse primary visual cortex. IScience 24, 102450.

Mumford, D. (1992). On the computational architecture of the neocortex II The role of cortico-cortical loops. Biol. Cybern. 66, 241–251.

Murakami, T., Matsui, T., and Ohki, K. (2017). Functional Segregation and Development of Mouse Higher Visual Areas. J. Neurosci. 37, 9424–9437.

Murakami, T., Matsui, T., Uemura, M., and Ohki, K. (2022). Modular strategy for development of the hierarchical visual network in mice. Nature 608, 578–585.

Pachitariu, M., Stringer, C., Dipoppa, M., Schröder, S., Rossi, L.F., Dalgleish, H., Carandini, M., and Harris, K.D. (2017). Suite2p: beyond 10,000 neurons with standard two-photon microscopy. BioRxiv 061507.

Pennartz, C.M.A., Dora, S., Muckli, L., and Lorteije, J.A.M. (2019). Towards a Unified View on Pathways and Functions of Neural Recurrent Processing. Trends Neurosci. 42, 589–603.

Pologruto, T.A., Sabatini, B.L., and Svoboda, K. (2003). ScanImage: flexible software for operating laser scanning microscopes. Biomed. Eng. Online 2, 13.

Puchades, M.A., Csucs, G., Ledergerber, D., Leergaard, T.B., and Bjaalie, J.G. (2019). Spatial registration of serial microscopic brain images to three-dimensional reference atlases with the QuickNII tool. PLoS One 14, 1–14.

Rao, R.P.N., and Ballard, D.H. (1999). Predictive coding in the visual cortex: a functional interpretation of some extra-classical receptive-field effects. Nat. Neurosci. 2, 79–87.

Rochefort, N.L., Narushima, M., Grienberger, C., Marandi, N., Hill, D.N., and Konnerth, A. (2011). Development of direction selectivity in mouse cortical neurons. Neuron 71, 425–432.

Rossi, L.F., Harris, K.D., and Carandini, M. (2020). Spatial connectivity matches direction selectivity in visual cortex. Nature 588, 648–652.

Schmidt, K.E., Goebel, R., Löwel, S., and Singer, W. (1997). The Perceptual Grouping Criterion of Colinearity is Reflected by Anisotropies of Connections in the Primary Visual Cortex. Eur. J. Neurosci. 9, 1083–1089.

Schneider, D.M., Sundararajan, J., and Mooney, R. (2018). A cortical filter that learns to suppress the acoustic consequences of movement. Nature 561, 391–395.

Sengpiel, F., Stawinski, P., and Bonhoeffer, T. (1999). Influence of experience on orientation maps in cat visual cortex. Nat. Neurosci. 2, 727–732.

Siegle, J.H., Jia, X., Durand, S., Gale, S., Bennett, C., Graddis, N., Heller, G., Ramirez, T.K., Choi, H., Luviano, J.A., et al. (2021). Survey of spiking in the mouse visual system reveals functional hierarchy. Nature 592, 86–92.

Sigman, M., Cecchi, G.A., Gilbert, C.D., and Magnasco, M.O. (2001). On a common circle: natural scenes and Gestalt rules. Proc. Natl. Acad. Sci. U. S. A. 98, 1935–1940.

Siu, C., Balsor, J., Merlin, S., Federer, F., and Angelucci, A. (2021). A direct interareal feedback-to-feedforward circuit in primate visual cortex. Nat. Commun. 12, 4911.

Stryker, M.P., Sherk, H., Leventhal, A.G., and Hirsch, H.V.B. (1978). Physiological consequences for the Cat’s visual cortex of effectively restricting early visual experience with oriented contours. J. Neurophysiol. 41, 896–909.

Summerfield, C., and Egner, T. (2009). Expectation (and attention) in visual cognition. Trends Cogn. Sci. 13, 403–409.

Suter, B.A., O’Connor, T., Iyer, V., Petreanu, L., Hooks, B.M., Kiritani, T., Svoboda, K., and Shepherd, G.M.G. (2010). Ephus: multipurpose data acquisition software for neuroscience experiments. Front. Neural Circuits 4, 100.

Tanaka, S., Tani, T., Ribot, J., and Yamazaki, T. (2007). Chronically mountable goggles for persistent exposure to single orientation. J. Neurosci. Methods 160, 206–214.

Trachtenberg, J.T. (2015). Competition, inhibition, and critical periods of cortical plasticity. Curr. Opin. Neurobiol. 35, 44–48.

Wang, Q., and Burkhalter, A. (2007). Area map of mouse visual cortex. J. Comp. 357, 339–357.

Wang, Q., Sporns, O., and Burkhalter, A. (2012). Network Analysis of Corticocortical Connections Reveals Ventral and Dorsal Processing Streams in Mouse Visual Cortex. J. Neurosci. 32, 4386–4399.

Yoshida, T., Ozawa, K., and Tanaka, S. (2012). Sensitivity Profile for Orientation Selectivity in the Visual Cortex of Goggle-Reared Mice. PLoS One 7, e40630.

Young, H., Belbut, B., Baeta, M., and Petreanu, L. (2021). Laminar-specific cortico-cortical loops in mouse visual cortex. Elife 10, 1–25.

Yu, Y., Stirman, J.N., Dorsett, C.R., and Smith, S.L. (2022). Selective representations of texture and motion in mouse higher visual areas. Curr. Biol. 32, 2810-2820.e5.

