## Supplemental Material for "Visual experience instructs the organization of cortical feedback inputs to primary visual cortex"

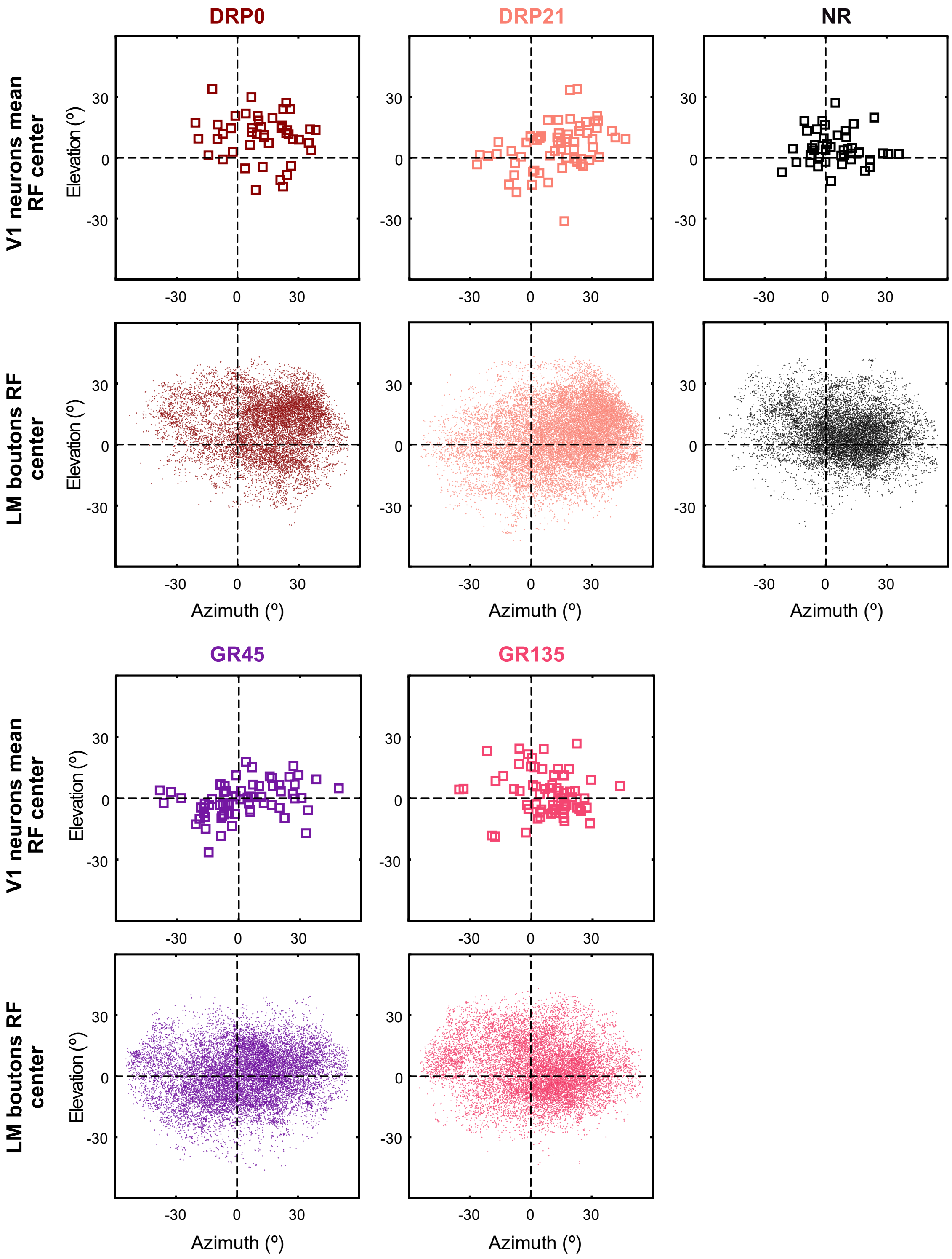


**Figure S1. Receptive field centers of V1 neurons and LM boutons for all experimental groups.**

**A**) Mean RF center of V1 neurons per imaging session (top) and RF center of all recorded boutons (bottom). (V1 neurons; DRP0 n=1560 from 45 imaging sessions in 6 animals; DRP21 n=1937 from 62 imaging sessions in 9 animals; NR n=1427 from 45 imaging sessions in 6 animals. LM boutons; DRP0 n=12597; DRP21 n=22517; NR n=13250)

**Figure S1**

**Dias, Rajan et al.**

**Figure S1. (legend continued)**

**B**) Same as **A** for the Goggle Reared groups. V1 neurons; GR45 n=2194 from 68 imaging sessions in 8 animals; GR135 n=1738 from 66 imaging sessions in 7 animals. LM boutons; GR45 n=16006; GR135 n=15049.

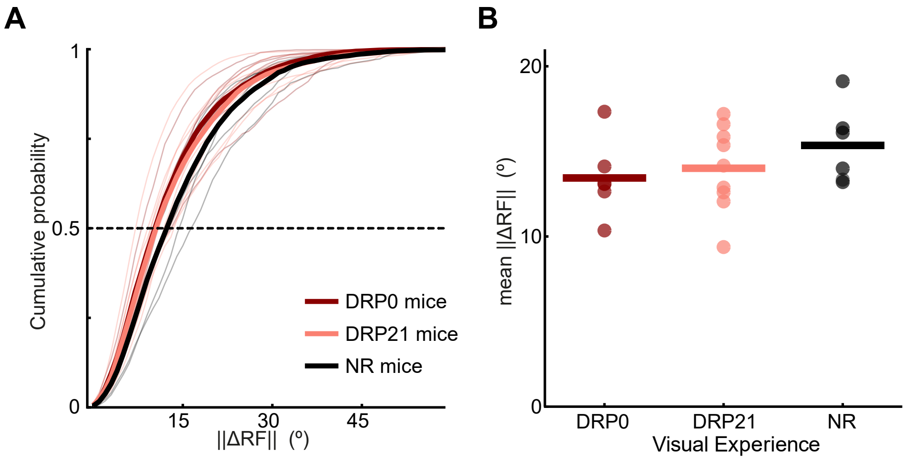


**Figure S2. ||ΔRF|| is unchanged in dark-reared mice.**

**A**) Cumulative probability distributions of ||ΔRF||. Thick lines, mean across animals for each group, thin lines, individual animals. DRP0, N=6; DRP21, N=9; NR, N=6.
**B**) Mean ||ΔRF|| of all boutons per animal (one-way ANOVA: $F_{2,18}$=1.03, p=0.38).

**Figure S2**

**Dias, Rajan et al.**

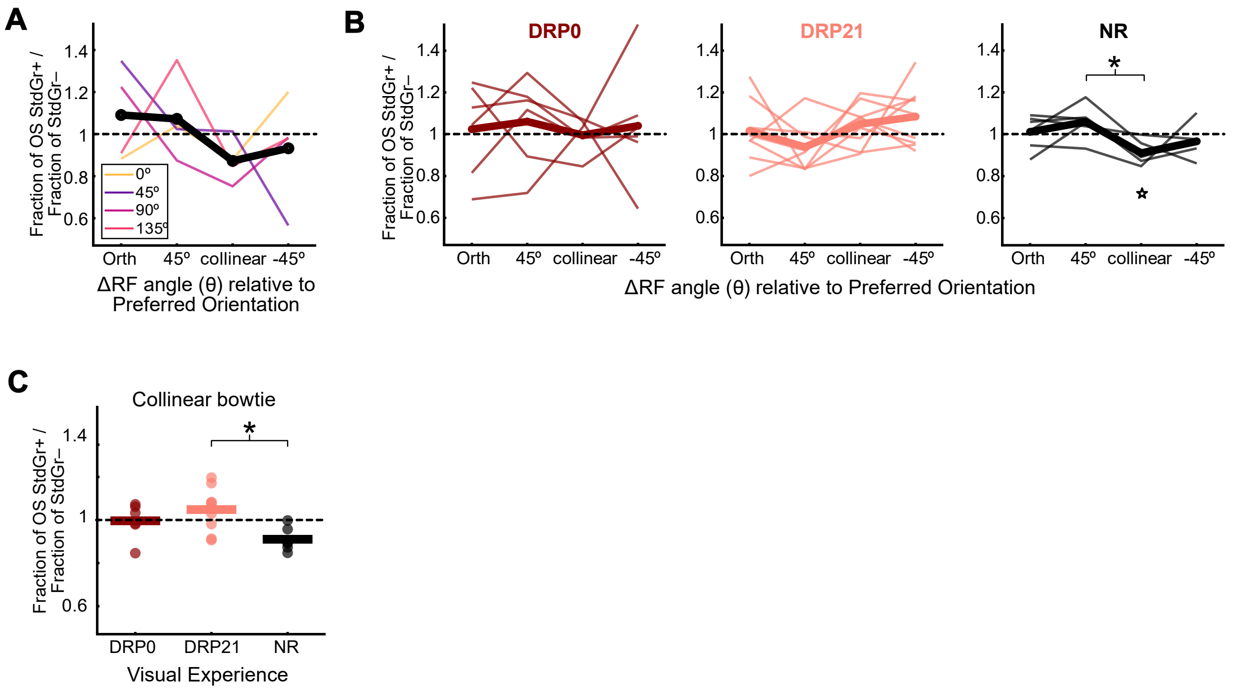


**Figure S3. When averaging across OS groups in NR mice, LM boutons are less likely to have ΔRF angle deviating collinearly to their preferred orientation.**

**A**) Colored lines, Fraction of OS boutons from a single example animal normalized to the non-grating selective population (StdGr–). Line colors indicate the different preferred orientations (one-way Repeated Measures ANOVA: $F_{3,3}$=0.8, p=0.53).
**B**) Fraction of OS StdGr+ boutons relative to the StdGr– population for the three visual experience groups (left) DRP0, (middle) DRP21, (right) NR (one-way Repeated Measures ANOVA: DRP0: $F_{5,3}$=0.08, p=0.97; DRP21: $F_{8,3}$=1.7, p=0.19; NR: $F_{5,3}$=3.8, p=0.033; Tukey-Kramer test: collinear vs. 45°, p=0.014; ☆t-test: collinear bowtie OS vs StdGr–, $t_5$=3.9, Bonferroni corrected p=0.046).
**C**) Fraction of OS StdGr+ boutons relative to the StdGr– population in the collinear bin across the three visual experience groups (one-way ANOVA: $F_{2,18}$=4.62, p=0.024; Tukey-Kramer test: DRP21 vs NR p=0.018).

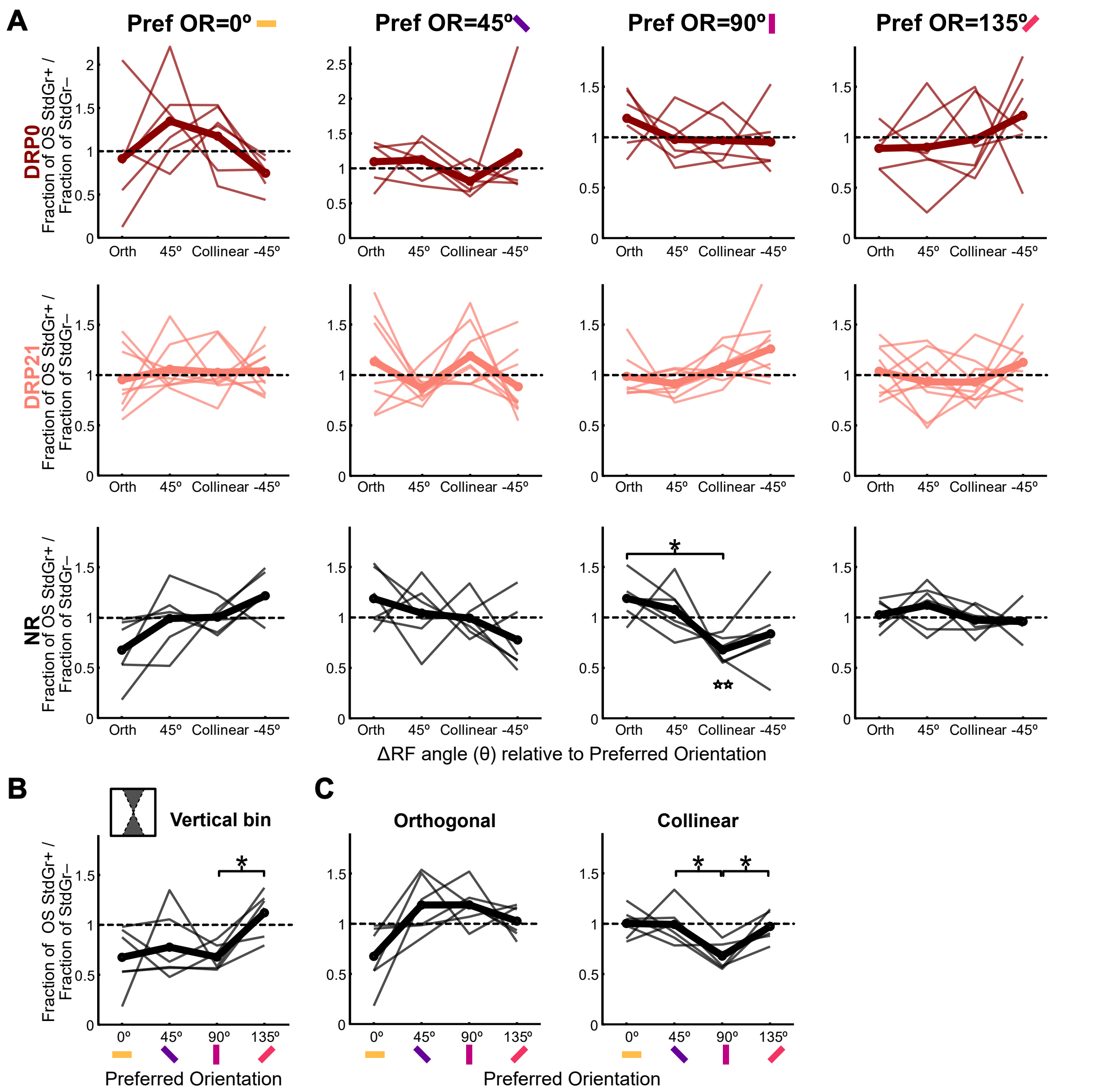


**Figure S4. Distribution of ΔRF deviation angle for LM boutons tuned to different orientations.**

**A**) Fraction of OS StdGr+ boutons relative to the StdGr– population for each OS population of LM boutons across the three visual experience groups (one-way Repeated Measures ANOVA; DRP0: 0°-preferring, $F_{5,3}$=1.56, p=0.22; 45°-preferring, $F_{5,3}$=0.72, p=0.56; 90°-preferring, $F_{5,3}$=0.76, p=0.53; 135°-preferring, $F_{5,3}$=0.72, p=0.56; DRP21: 0°-preferring, $F_{8,3}$=0.21, p=0.89; 45°-preferring, $F_{8,3}$=1.92, p=0.15; 90°-preferring, $F_{8,3}$=2.98, p=0.051; 135°-preferring, $F_{8,3}$=0.85, p=0.48; NR: 0°-preferring, $F_{5,3}$=3.50, p=0.042; 45°-preferring, $F_{5,3}$=1.56, p=0.24; 90°-preferring, $F_{5,3}$=3.71, p=0.035; 135°-preferring, $F_{5,3}$=0.81, p=0.51; *Tukey Kramer test: 90°-preferring: collinear vs orthogonal, p=0.016, ☆☆ t-test: 90°-preferring vs. StdGr– , Bonferroni corrected $t_5$=6.01, p=0.007).

### Figure S4

**Dias, Rajan et al.**

**Figure S4. (legend continued)**

**B**) Fraction of OS StdGr+ boutons relative to the StdGr– population in the vertical bowtie across the different preferred orientations for NR mice (vertical bowtie: one-way Repeated Measures ANOVA: $F_{5,3}$=3.56, p=0.04; Tukey-Kramer test, 90° vs. 135°, p=0.024).

**C**) Fraction of OS StdGr+ boutons relative to the StdGr– population in the orthogonal and collinear bowties across the different preferred orientations for NR mice (one-way Repeated Measures ANOVA; orthogonal bowtie bin, $F_{5,3}$=4.92, p=0.014; Collinear bowtie bin, $F_{5,3}$=5.70, p=0.0083; Tukey-Kramer test, 90°-preferring vs. 45°-preferring, p=0.028; 90°-preferring vs. 135°-preferring, p=0.042).
